# Endoglycan plays a role in axon guidance and neuronal migration by negatively regulating cell-cell adhesion

**DOI:** 10.1101/425207

**Authors:** Thomas Baeriswyl, Alexandre Dumoulin, Martina Schaettin, Georgia Tsapara, Vera Niederkofler, Denise Helbling, Evelyn Avilés, Jeannine A. Frei, Nicole H. Wilson, Matthias Gesemann, Esther T. Stoeckli

## Abstract

Cell migration and axon guidance are important steps in the formation of neural circuits. Both steps depend on the interactions between cell surface receptors and molecules on cells along the pathway. In addition to cell-cell adhesion, these molecular interactions provide guidance information. The fine-tuning of cell-cell adhesion is an important aspect of cell migration, axon guidance, and synapse formation.

Here, we show that Endoglycan, a sialomucin, plays a role in axon guidance and cell migration in the central nervous system. In the absence of Endoglycan, commissural axons failed to properly navigate the midline of the spinal cord. In the developing cerebellum, a lack of Endoglycan prevented migration of Purkinje cells and resulted in a stunted growth of the cerebellar lobes. Taken together, these results support the hypothesis that Endoglycan acts as a ‘lubricant’, a negative regulator of cell-cell adhesion, in both commissural axon guidance and Purkinje cell migration.

## INTRODUCTION

Cell migration and axonal pathfinding are crucial aspects of neural development. Neurons are born in proliferative zones from where they migrate to their final destination. After their arrival, they send out axons that have to navigate through the tissue to find the target cells with which they establish synaptic contacts. Intuitively, it is clear that the same cues provided by the environment can guide both cells and axons to their target. Although we know relatively little about guidance cues for cells compared to guidance cues for axons, both processes are dependent on proper cell-cell contacts (Gomez and Letourneau, 2014; Short et al., 2016). One of the best-studied model systems for axon guidance are the commissural neurons located in the dorsolateral spinal cord (de Ramon Francàs et al., 2017; Stoeckli, 2017 and 2018). These neurons send out their axons toward the ventral midline under the influence of the roof plate-derived repellents BMP7 (Augsburger et al., 1999) and Draxin (Islam et al., 2009). At the same time, axons are attracted to the floor plate, their intermediate target, by Netrin (for a review on Netrin function, see Boyer and Gupton, 2018), VEGF (Ruiz de Almodóvar et al., 2011), and Shh (Yam et al., 2009 and 2012). At the floor-plate border, commissural axons require the short-range guidance cues Contactin2 (aka Axonin-1) and NrCAM to enter the midline area (Stoeckli and Landmesser, 1995; Stoeckli et al., 1997; Fitzli et al., 2000; Pekarik et al., 2003). Slits and their receptors, the Robos, were shown to be required as negative signals involved in pushing axons out of the midline area (Long et al., 2004; Blockus and Chédotal, 2016). Members of the Semaphorin family are also involved in midline crossing either as negative signals mediated by Neuropilin-2 (Zou et al., 2000; Parra and Zou, 2010; Nawabi et al., 2010; Charoy et al., 2012) or as receptors for floor-plate derived PlexinA2 (Andermatt et al., 2014a). Once commissural axons exit the floor-plate area, they turn rostrally along the longitudinal axis of the spinal cord. Morphogens of the Wnt family (Lyuksyutova et al., 2003; Domanitskaya et al., 2010; Avilés and Stoeckli, 2016) and Shh (Bourikas et al., 2005; Wilson and Stoeckli, 2013) were identified as guidance cues directing post-crossing commissural axons rostrally. In the same screen that resulted in the discovery of Shh as a repellent for post-crossing commissural axons (Bourikas et al., 2005), we found another candidate that interfered with the rostral turn of post-crossing commissural axons. This candidate gene was identified as *Endoglycan*.

Endoglycan is a member of the CD34 family of sialomucins (Nielsen and McNagny, 2008; Sassetti et al., 2000; Furness and McNagny, 2006). The family includes CD34, Podocalyxin (also known as Thrombomucin, PCLP-1, MEP21, or gp135), and Endoglycan (also known as Podocalyxin-like 2). They are single-pass transmembrane proteins with highly conserved transmembrane and cytoplasmic domains. A C-terminal PDZ recognition site is found in all three family members (Furness and McNagny, 2006; Nielsen and McNagny, 2008). The hallmark of sialomucins is their bulky extracellular domain that is negatively charged due to extensive N- and O-glycosylation. Despite the fact that CD34 was identified more than 20 years ago, very little is known about its function. It has been widely used as a marker for hematopoietic stem cells and precursors. Similarly, Podocalyxin is expressed on hematopoietic stem and precursor cells. In contrast to CD34, Podocalyxin was found in podocytes of the developing kidney (Kerjaschki et al., 1984; Doyonnas et al., 2005; Furness and McNagny, 2006). In the absence of Podocalyxin, podocytes do not differentiate, resulting in kidney failure and thus perinatal lethality in mice (Doyonnas et al., 2001). *Podocalyxin*, but not *CD34*, is expressed widely in the developing and mature mouse brain (Vitureira et al., 2005; Vitureira et al., 2010). Podocalyxin was shown to induce microvilli and regulate cell-cell adhesion via its binding to the NHERF (Na^+^/H^+^ exchanger regulatory factor) family of adaptor proteins that link Podocalyxin to the actin cytoskeleton (Nielsen et al., 2007; Nielsen and McNagny, 2008; Nielsen and McNagny, 2009). Like *Podocalyxin*, *Endoglycan* is expressed in the brain and in the kidney. Only low levels were found in hematopoietic tissues (Sassetti et al., 2000). To date, nothing is known about the function of Endoglycan.

Based on its temporal and spatial expression pattern, we first analyzed the function of Endoglycan in the embryonic chicken spinal cord. In the absence of Endoglycan commissural axons failed to turn rostrally upon floor-plate exit. Occasionally, they were observed to turn already inside the floor-plate area. Furthermore, the trajectory of commissural axons in the midline area was tortuous in embryos lacking Endoglycan, but straight in control embryos. Live imaging data of dI1 axons crossing the floor plate confirmed changes in axon – floor plate interaction. In addition, axons were growing more slowly after silencing *Endoglycan* and faster after overexpression of *Endoglycan*.

In the cerebellum, *Endoglycan* expression is restricted to migrating Purkinje cells. The absence of Endoglycan resulted in the failure of Purkinje cells to migrate properly from the ventricular zone to their destination in the periphery of the cerebellum, where they normally form the typical Purkinje cell layer. This in turn resulted in a decrease in granule cell proliferation and in the stunted growth of the cerebellar folds.

Taken together, our results from in vitro and in vivo studies along with live imaging observations are consistent with a role for Endoglycan as a ‘lubricant’, a negative regulator of cell-cell contact and therefore modulator of molecular interactions affecting both axon guidance and cell migration.

## RESULTS

### Endoglycan was identified as a candidate guidance cue for commissural axons

In a subtractive hybridization screen, we identified differentially expressed floor-plate genes as candidate guidance cues directing axons from dorsolateral commissural neurons (dI1 neurons) along the longitudinal axis after midline crossing (Bourikas et al., 2005; see Methods). Candidates with an expression pattern that was compatible with a role in commissural axon navigation at the midline were selected for functional analysis using in ovo RNAi (Pekarik et al., 2003; Wilson and Stoeckli, 2011). One of these candidates that interfered with the correct rostral turning of commissural axons after midline crossing turned out to be *Endoglycan*, a member of the CD34 family of sialomucins.

CD34 family members share a common domain organization that consists of a mucin-like domain followed by a cysteine-containing globular domain, a membrane associated stalk region, a transmembrane spanning domain and the cytoplasmic domain (Supplementary Figure 1; Sassetti et al., 2000; Furness and McNagny, 2006; Nielsen and McNagny, 2008). With the exception of the mucin-like domain at the N-terminus, the conservation between species orthologues of CD34, Endoglycan and Podocalyxin is in the range of 80%, but drops below 40% within the mucin domain. However, homologies of these paralogous proteins within the same species are generally only in the range of 40% (Supplementary Figure 1), demonstrating that, while they might share a similar overall structure, the structure can be built by quite diverse primary amino acid sequences.

*Endoglycan* was expressed mainly in the nervous system during development, as levels in non-neuronal tissues were much lower (Supplementary Figure 2). In the neural tube, *Endoglycan* was expressed ubiquitously including floor-plate cells at HH21 (Hamburger and Hamilton stage 21; Hamburger and Hamilton, 1951; Figure 1). By HH25, expression was still found throughout the neural tube withhigher levels detected in dorsal neurons (including dI1 neurons) and motoneurons. *Endoglycan* expression was also maintained in the floor plate (Figure 1B). For functional analysis, dsRNA was produced from the *Endoglycan* cDNA fragment obtained from a screen and used for in ovo electroporation of the spinal cord at HH18 (Figure 1D). The analysis of commissural axons’ trajectories at HH26 by DiI tracing in „open-book“ preparations (Figure 1E; quantified in Figure 1L) revealed either failure to turn or erroneous caudal turns along the contralateral floor-plate border in embryos lacking Endoglycan in the floor plate (Figure 1G), in only the dorsal spinal cord (Figure 1J), or in one half of the spinal cord including the floor plate (Figure 1H,I). Furthermore, axons were occasionally found to turn prematurely either before midline crossing or within the floor-plate area. Detailed analysis of the axonal morphology in the floor-plate area revealed a tortuous, ‘corkscrew’-like trajectory in embryos lacking Endoglycan in dI1 neurons and the floor plate (Figure 1I), whereas axons crossed the midline in a straight trajectory in untreated control embryos (Figure 1F) and in embryos injected and electroporated with dsRNA derived from either CD34 (not shown) or Podocalyxin (Figure 1K).

**Figure 1.**
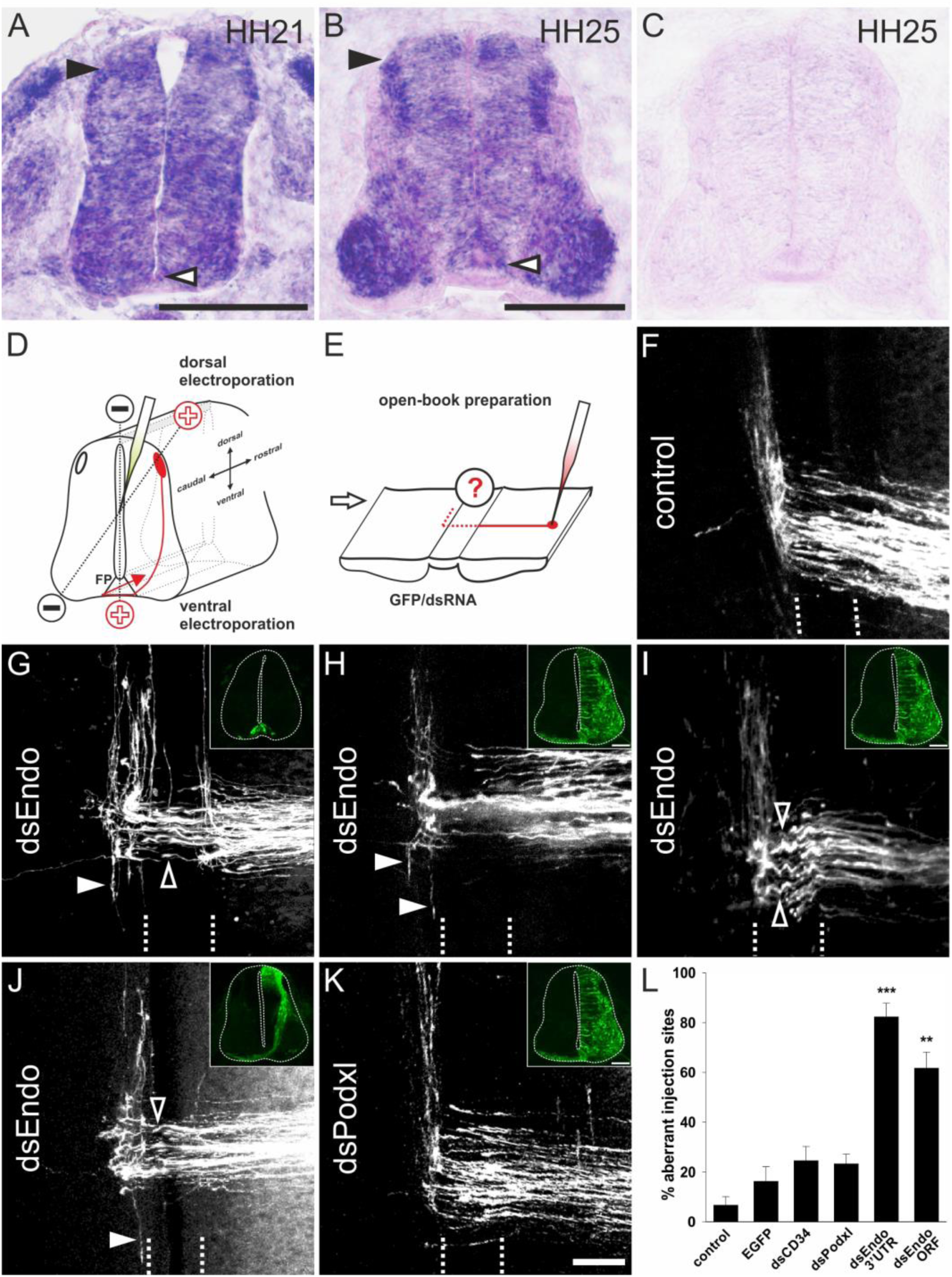
Endoglycan is required for correct turning of post-crossing commissural axons. (A,B) *Endoglycan* is expressed in the developing neural tube during commissural axon guidance. *Endoglycan* is expressed throughout the neural tube at HH21, including the floor plate (white arrowhead) (A). (B) At HH25, *Endoglycan* is still found in most cells of the spinal cord. High levels are found in motoneurons and interneurons, including the dorsal dI1 neurons (black arrowhead), and in the floor plate (white arrowhead). No staining was found when hybridization was carried out with a sense probe (C). Commissural axon pathfinding was analyzed in “open-book” preparations (D,E; see Methods for details). The positions of the electrodes for dorsal and ventral electroporation are indicated (D). In control embryos at HH26, commissural axons have crossed the floor plate and turned rostrally along the contralateral floor-plate border (F). In contrast, after downregulation of Endoglycan (G-J) commissural axons failed to turn along the contralateral floor-plate border or they turned randomly either rostrally or caudally (arrowheads in G-J). Occasionally axons were turning already inside the floor plate (open arrowhead in G). A closer look at the morphology of the axons in the floor plate revealed their tortuous, ‘corkscrew-like’ trajectory across the midline at many DiI injection sites (open arrowheads in I). To knockdown *Endoglycan* either in the floor plate or in commissural neurons, the ventral or dorsal spinal cord was targeted as indicated in (D) (see inserts in G and J, respectively). Phenotypes were the same as those observed after targeting one half of the spinal cord including the floor plate (H,I). Pathfinding was normal in embryos electroporated with dsRNA derived from *Podocalyxin* (K). The quantification of injection sites with pathfinding errors after targeting the floor plate or one half of the spinal cord is shown in (L). Pathfinding errors were seen only at 6.7±3.4% of the injection sites in untreated control embryos (n=10 embryos, 45 injection sites). In control embryos injected and electroporated with the *EGFP* plasmid alone, pathfinding errors were found at 16.2±6% of the injection sites (n=17 embryos, 92 injection sites). Injection and electroporation of dsRNA derived from *CD34* (24.6±5.8%, n=8 embryos, 80 injection sites) and *Podocalyxin* (23.3±3.9%, n=17 embryos, 147 injection sites) did not affect midline crossing and turning behavior of commissural axons. In contrast, 82.3±5.6% (n=11 embryos, 65 sites) and 61.7±6.4% (n=18, 161 sites) of the injection sites in embryos injected with dsRNA derived from the 3’-UTR or the ORF of *Endoglycan*, respectively, showed aberrant pathfinding of commissural axons. P values ***<0.001 and **<0.01, compared to EGFP-injected control groups. The two groups electroporated with dsRNA derived from *Endoglycan* were not different from each other. Values represent average percentage of DiI injection sites per embryo with aberrant axonal navigation ± standard error of the mean. Bar: 50 µm.

To demonstrate specificity of Endoglycan downregulation and to verify that the phenotype was not due to an off-target effect, we used three non-overlapping cDNA fragments to produce dsRNA. All fragments resulted in the same phenotypes. Downregulation of Endoglycan with dsRNA derived from the ORF resulted in 61.7±6.4% injection sites with aberrant axon guidance. The effect on axon guidance was also seen with dsRNA derived from the 3’UTR, with 82.3±5.6% of the injection sites with aberrant axon guidance (Figure 1L). In contrast, aberrant axonal pathfinding was seen only at 6.7±3.4% of the injection sites in untreated control embryos. Values were 16.2±6.0 for EGFP-expressing control embryos, 24.6±5.8% for embryos transfected with dsRNA derived from *CD34*, and 23.3±3.9% for embryos transfected with dsRNA derived from *Podocalyxin*. Thus, silencing either *CD34* or *Podocalyxin* did not interfere with correct navigation of axons at the midline. Because both of them were expressed in the developing spinal cord (Supplementary Figure 3), these results further support the specificity of the observed effects of *Endoglycan* silencing.

### Lack of Endoglycan affects the morphology of the floor plate only after dI1 axons have crossed the midline

Because the hallmark of sialomucins is their bulky, negatively charged extracellular domain with extensive glycosylation, a role as regulators of cell-cell adhesion has been postulated (Vitureira et al., 2010; Takeda et al., 2000; Nielsen and McNagny, 2008 and 2009). This together with our observation that commissural axons have a „corkscrew“-like phenotype in the midline area in Endoglycan-deficient embryos prompted us to analyze the morphology of the floor plate. Sections were taken from the lumbar level of the spinal cord at HH26 from control-treated and experimental embryos and stained for HNF3β/FoxA2 to label floor-plate cells, and for Contactin2 (aka Axonin-1) to label commissural axons (Figure 2). In untreated (Figure 2A-C) and control-treated embryos (Figure 2D-F), HNF3β/FoxA2-positive cells were aligned to form the characteristic triangular shape of the floor plate. In particular, the ventral border of the floor plate, where commissural axons traverse the midline was smooth, because all floor-plate cells were precisely aligned (Figure 2A,D). In contrast, floor-plate cells were no longer aligned to form a smooth ventral border in embryos lacking Endoglycan (Figure 2G,J). On the one hand, floor-plate cells were found dislocated into the commissure formed by the Contactin2-positive axons (arrowheads in Figure 2I,L). On the other hand, the floor plate appeared to have gaps in embryos lacking Endoglycan. In addition, the floor-plate width was significantly narrower in embryos lacking Endoglycan in comparison to age-matched controls (Figure 2S,T). These changes in floor-plate morphology were not due to differences in cell differentiation or patterning (Supplementary Figure 4). Furthermore, we can exclude cell death as a contributor to the changes in the floor plate, as we found no Cleaved Caspase-3-positive floor-plate cells in any of the conditions.

**Figure 2.**
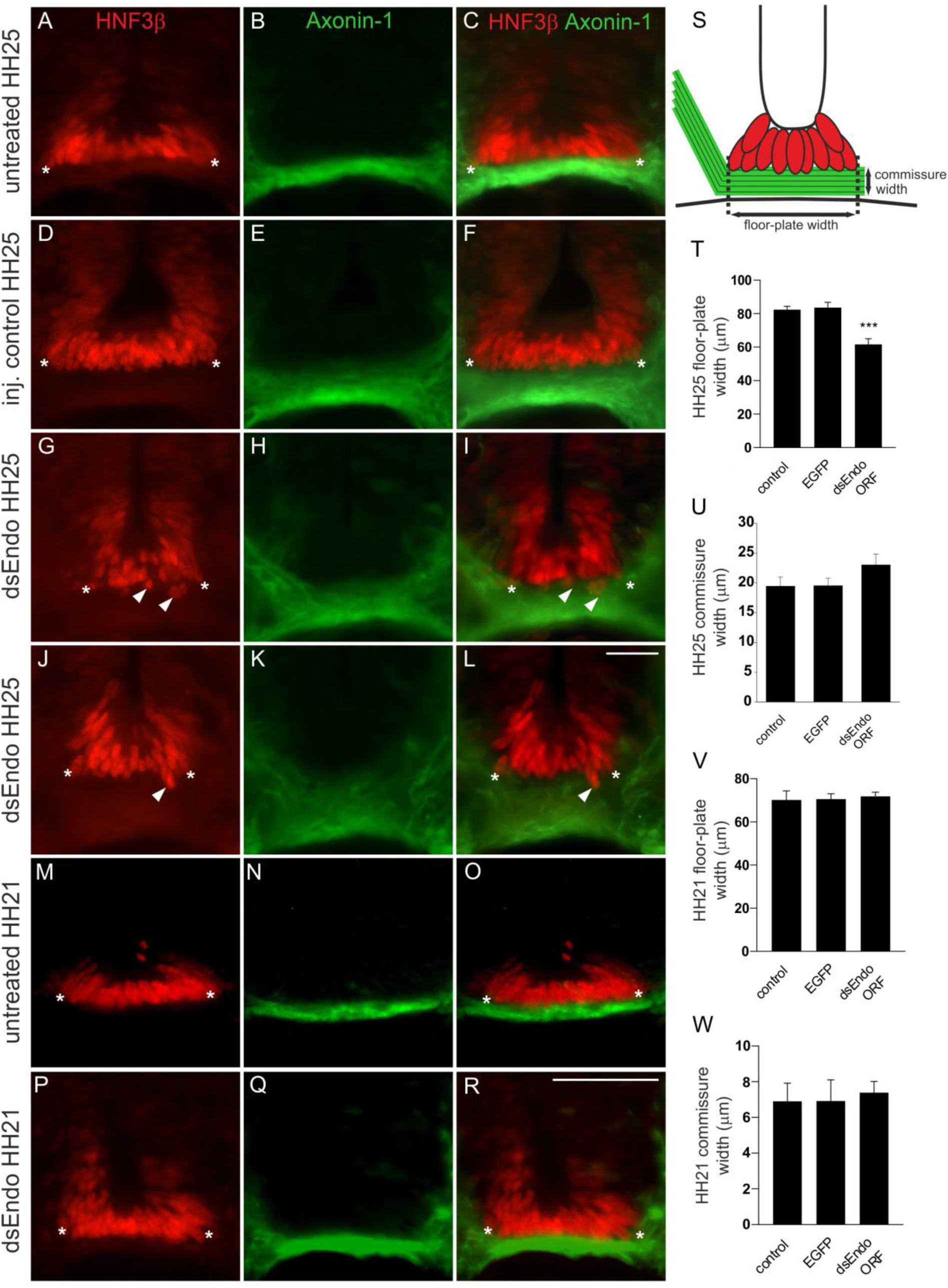
After downregulation of Endoglycan the floor-plate morphology is compromised only after axonal midline crossing. In untreated (A-C) and control-treated embryos (D-F) the floor plate is of triangular shape with floor-plate cells precisely aligned at the ventral border. There is no overlap between the floor plate (visualized by HNF3β staining; red) and the commissure (visualized by anti-Axonin1 staining; green). The shape of the floor plate is no longer triangular in embryos lacking Endoglycan (G-L). The floor-plate cells are not aligned ventrally (arrowheads in G, I, J, and L) and the floor plate appears to have gaps. This change in morphology is only seen at HH25, when midline crossing is completed. When the floor-plate morphology was analyzed at HH21, there was no difference between control (M-O) and experimental embryos electroporated with dsRNA derived from *Endoglycan* (P-R). Note that some more ventral commissural axon populations have crossed the floor plate at this stage. But overall, the number of axons that form the commissure at HH21 is still very small. The width of the floor plate (indicated by asterisks) was measured (S,T). There was no significant difference in spinal cord width at HH25 (400.2±54.5 µm in untreated controls, 438.2±30.3 µm in EGFP-expressing controls, and 394±12.0 µm in dsEndo embryos), but floor plates were significantly narrower in embryos lacking Endoglycan (T; 61.6±3.4 µm; n=7 embryos; p<0.001) compared to untreated (82.4±2.0 µm; n=6 embryos) and *EGFP*-injected control embryos (83.6±3.2 µm; n=6 embryos). The commissure had a tendency to be wider in experimental compared to control embryos but the effect was not statistically significant (U). The width of the floor plate was not different between groups when measured at HH21 (V) with 70.1±4.2 µm (n=3) for untreated and 70.6±2.5 µm for EGFP controls (n=4), compared to 71.8±2.0 µm for experimental embryos (n=3). No difference was seen in the width of the commissure. Bar: 50 µm. ANOVA with Bonferroni correction was used for statistical analyses.

When embryos lacking Endoglycan were analyzed at HH21 (Figure 2P-R), that is at a time point when dI1 axons have reached but not yet crossed the floor plate, the morphology and the width of the floor plate were not different from controls (Figure 2M-O). In contrast to measurements of floor-plate width at HH25 (Figure 2T), the values for experimental and control embryos were not different at HH21 (Figure 2V). Thus, we concluded that the absence of Endoglycan did not affect primarily cell-cell adhesion between floor-plate cells. Rather the altered floor-plate morphology appeared to be an indirect effect of changes in axon to floor-plate adhesion. We ruled out an effect of the perturbation of Endoglycan levels on the expression of known guidance cues of dI1 axons, such as Contactin2 (Axonin-1) or NrCAM (Supplementary Figure 5). Similarly, we did not find changes in the expression of Shh and Wnt5a, morphogens that are known to direct post-crossing dI1 axons rostrally (Supplementary Figure 6; Bourikas et al., 2005; Lyuksyutova et al., 2003).

An alternative way of demonstrating the requirement for Endoglycan in both floor plate and commissural axons were rescue experiments (Figure 3). We used dsRNA derived from the 3’-UTR and expressed the ORF of *Endoglycan* either under control of the Math1 enhancer (expression only in dI1 neurons) or the Hoxa1 enhancer for floor-plate specific expression. Because expression of these plasmids in control embryos (overexpression) resulted in aberrant behavior of axons at the floor plate, we used three different concentrations of plasmid for our rescue experiments and obtained a dose-dependent effect on axon guidance. Expression of high doses of *Endoglycan* was never able to rescue the axon guidance phenotype. However, axon guidance was not different from control embryos after transfection of dI1 neurons with a low concentration or floor-plate cells with a medium concentration of the *Endoglycan* ORF (Figure 3B; Table 1). Interestingly, the source of Endoglycan did not matter but the amount of Endoglycan did. These findings were consistent with the idea that Endoglycan was regulating adhesion, as both too much but also too little adhesion would be a problem for axonal navigation. Furthermore, these results suggested that Endoglycan did not act as a receptor.

**Figure 3.**
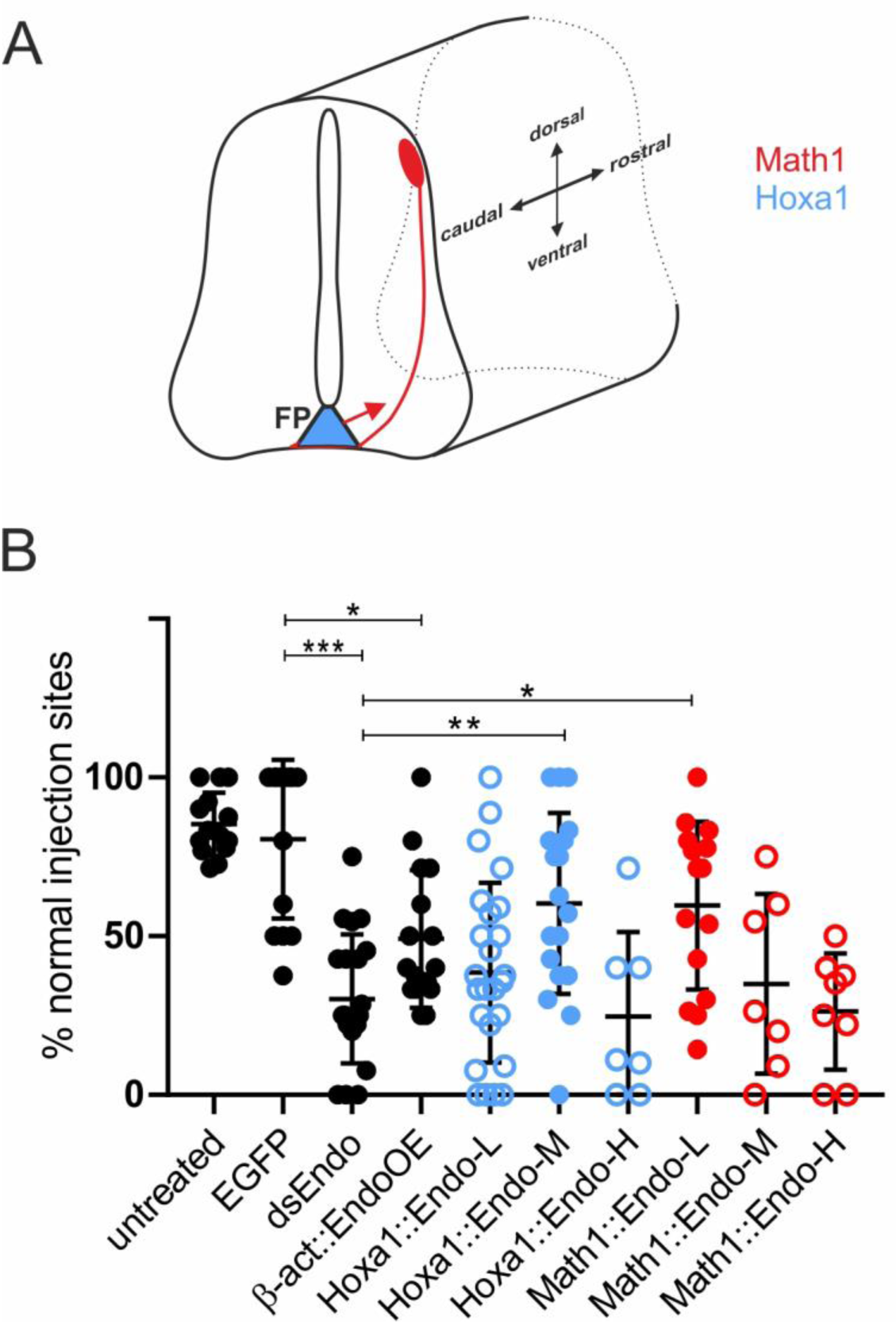
Too much or too little Endoglycan causes aberrant axon guidance. Because silencing *Endoglycan* either in commissural neurons or in the floor plate caused the same type of axon guidance defects, we wanted to test the idea that the presence of an adequate amount, but not the source of Endoglycan was important. We therefore downregulated Endoglycan by transfection of dsRNA derived from the 3’UTR of *Endoglycan* into one half of the spinal cord. We then tried to rescue the aberrant axon guidance phenotype by co-electroporation of the *Endoglycan* ORF specifically in dI1 neurons (using the Math1 enhancer, red) or in the floor plate (using the Hoxa1 enhancer; blue, A). The rescue constructs were used at a concentration of 150 (L = low), 300 (M = medium), and 750 (H = high) ng/µl, respectively. In both cases, rescue was only possible with one concentration: the medium concentration of the Endoglycan plasmid driven by the Hoxa1 promoter and the low concentration of the plasmid driven by the Math1 promoter. The lowest concentration of the Hoxa1-driven construct and the two higher concentrations of the Math1-driven constructs were not able to rescue the aberrant phenotype. Note that the amounts of Endoglycan cannot be compared between the Math1- and the Hoxa1 enhancers, as they differ in their potency to drive expression. However, we can conclude a response in a dose-dependent manner in both cases. Statistical analysis by one-way ANOVA: *p<0.05, **p<0.01, ***p<0.001. See Table 1 for quantification.

**Table 1.** The source of Endoglycan does not matter. Concomitant expression of Endoglycan could rescue the aberrant axon guidance phenotype induced by the downregulation of Endoglycan throughout the spinal cord. It did not matter whether Endoglycan was expressed under the Hoxa1 enhancer for specific expression in floor-plate cells, or under the Math1 enhancer for specific expression in dI1 neurons. However, the rescue effect was dose-dependent. Too little, or too much Endoglycan was inducing axon guidance defects. For rescue, *Endoglycan* cDNA under the control of the Hoxa1 enhancer (Hoxa1::Endo) or the Math1 enhancer (Math1::Endo) were injected at 150 ng/µl (low, L), 300 ng/µl (medium, M), or 750 ng/µl (high, H). The number of embryos and the number of DiI injection sites analyzed per group are indicated. The average % of injection sites with normal axon guidance phenotypes and the P-value for the comparison between the respective group and the control-treated (EGFP-expressing) group are given.

**The source of Endoglycan does not matter**
| Treatment | No of embryos | No of inj. sites | % inj sites<br>normal PT | P-value |
| --- | --- | --- | --- | --- |
| untreated | 14 | 111 | 85.3 ± 2.6 | 0.999 |
| GFP | 14 | 85 | 80.5± 6.7 | 1 |
| dsEndo | 15 | 111 | 30.2±5.3 | <0.0001 |
| β-actin::EndoOE | 16 | 91 | 49.1±2.2 | 0.0184 |
| Hoxa1::Endo-L | 25 | 234 | 38.5±5.7 | <0.0001 |
| Hoxa1::Endo-M | 18 | 136 | 60.3±6.7 | 0.3403 |
| Hoxa1::Endo-H | 6 | 64 | 24.6±10.9 | <0.0001 |
| Math1::Endo-L | 15 | 157 | 59.6±6.8 | 0.3515 |
| Math1::Endo-M | 7 | 86 | 35±10.7 | 0.0032 |
| Math1::Endo-H | 8 | 86 | 26.3±6.5 | <0.0001 |

### Endoglycan is a negative regulator of cell adhesion

The observation that downregulation of Endoglycan seemed to increase the adhesion between commissural axons and floor-plate cells, together with the knowledge about its molecular features, led us to hypothesize that Endoglycan might act as a negative regulator of cell-cell adhesion. As a first test, we counted the number of motoneurons that adhered to a carpet of HEK cells stably expressing Endoglycan compared to control HEK cells (Figure 4). Expression of Endoglycan interfered strongly with motoneuron adhesion, as the number of cells was almost three-fold higher on control HEK cells. The anti-adhesive properties of Endoglycan were explained by its post-translational modification (Supplementary Figure 7). Enzymatic removal of sialic acid by Neuraminidase or of O-linked glycans by O-glycosidase abolished the anti-adhesive properties of Endoglycan expressed in HEK cells.

**Figure 4.**
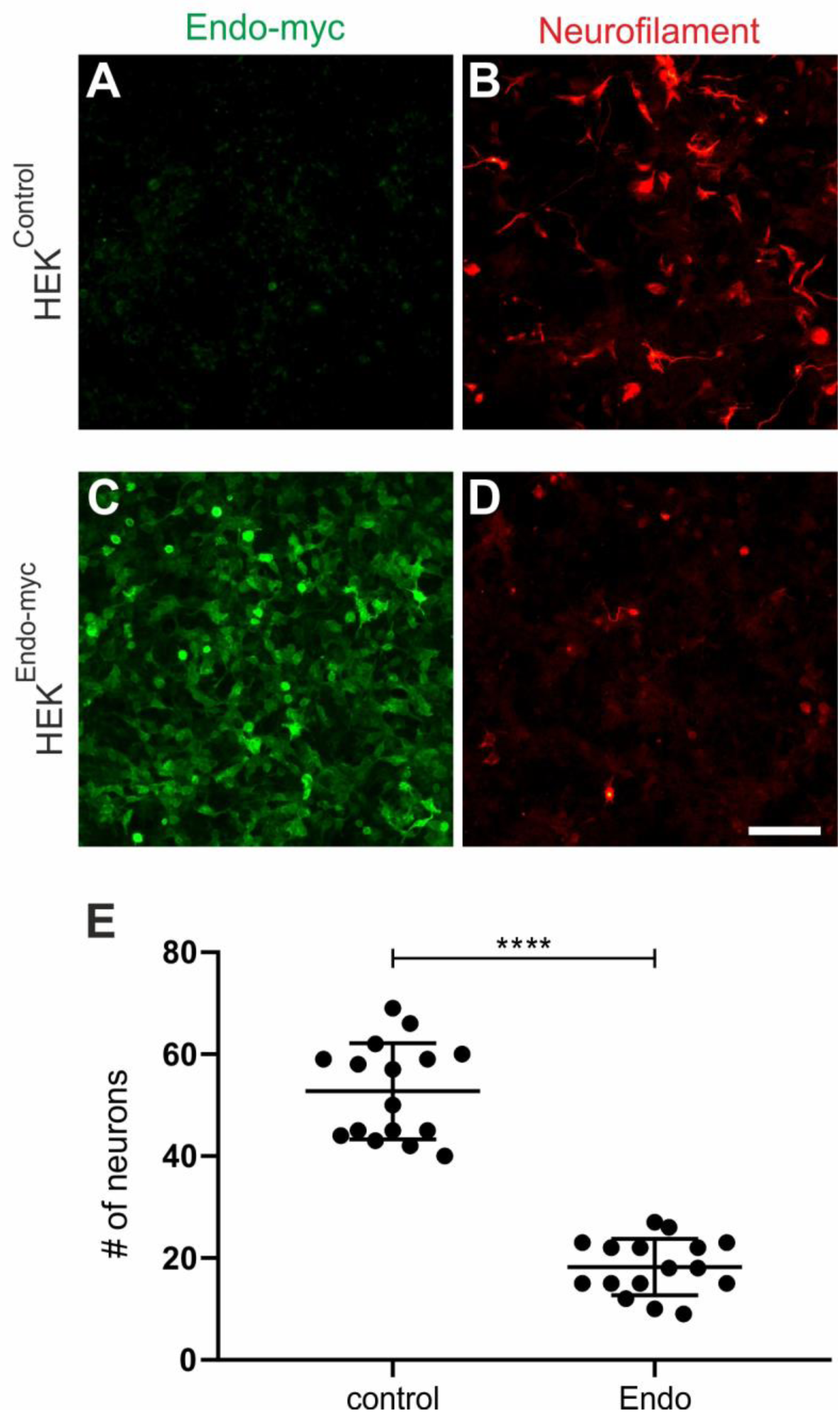
Endoglycan expression reduces cell adhesion in vitro. Control HEK cells (A) or HEK cells stably expressing human Endoglycan (C) were used as carpet for dissociated motoneurons dissected from HH26 chicken embryos (B,D). Neurons were allowed to attach for 40 hours. Staining for Neurofilament revealed a pronounced decrease in the number of motoneurons on HEK cells expressing Endoglycan (D) compared to control HEK cells (B). On control HEK cells (A), we found 52.8±2.4 motoneurons per view field. On a carpet of HEK cells expressing Endoglycan only 18.3±1.4 motoneurons were counted. Similar results were obtained in five independent experiments. Bar: 100 μm. ****p<0.0001, unpaired t test, two-tailed.

In addition, we tested our hypothesis that Endoglycan was a negative regulator of adhesion by manipulating the balance of adhesion between commissural axons and the floor plate. We had previously used a similar approach to demonstrate a role of RabGDI in Robo trafficking (Philipp et al., 2012). Commissural axons cross the midline because of the positive signals provided by the interaction of floor-plate NrCAM with growth cone Contactin2 (Stoeckli and Landmesser, 1995; Stoeckli et al., 1997; Fitzli et al., 2000). In the absence of NrCAM or Contactin2, commissural axons fail to enter the floor plate and turn into the longitudinal axis prematurely along the ipsilateral floor-plate border. The positive signal derived from the Contactin2/NrCAM interaction depends on sufficient contact between growth cone and floor-plate cells. Thus, we hypothesized that the failure to detect the positive signal due to lower NrCAM levels on the floor-plate cells could be counteracted by a forced increase in growth cone-floor plate contact. We reasoned that the concomitant downregulation of NrCAM and Endoglycan would rescue the NrCAM phenotype, because the decrease in adhesion due to lower NrCAM, resulting in the failure of commissural axons to enter the floor plate, would be counteracted by an increase in adhesion in the absence of Endoglycan. This is indeed what we observed (Figure 5). As found previously (Stoeckli and Landmesser, 1995; Pekarik et al., 2003), axons were frequently turning prematurely along the ipsilateral floor-plate border in the absence of NrCAM (Figure 5A). In accordance with our hypothesis, ipsilateral turns were reduced to control levels when NrCAM and Endoglycan were downregulated concomitantly (Figure 5B,G). The rescue of the NrCAM phenotype was only seen for Endoglycan, as concomitant downregulation of NrCAM with Podocalyxin or CD34 had no effect on ipsilateral turns (Figure 5).

**Figure 5.**
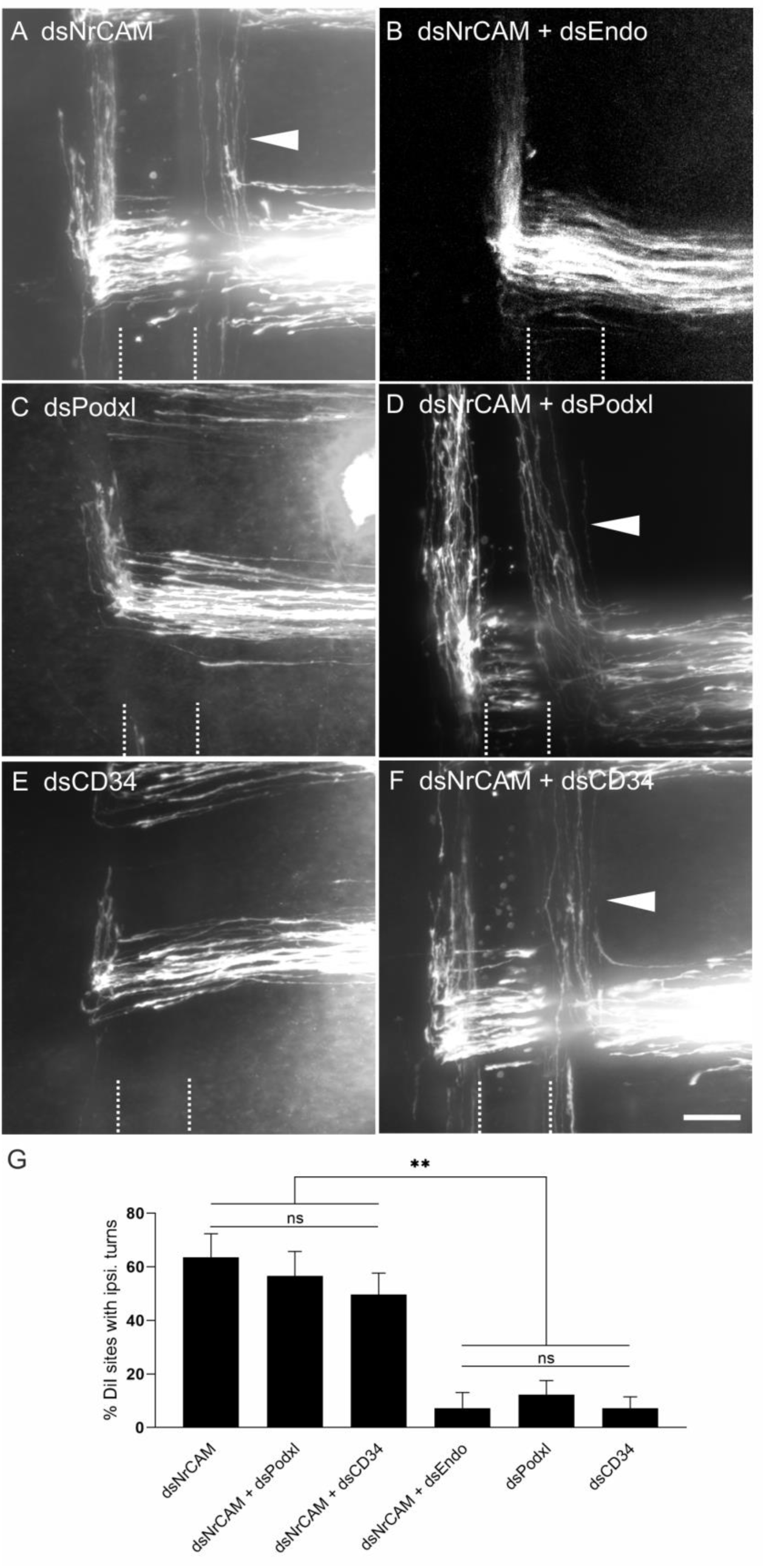
Downregulation of Endoglycan but not its family members rescues the axon guidance phenotype induced by downregulation of NrCAM. The perturbation of axon/floor-plate contact by downregulation of NrCAM resulted in the failure of commissural axons to enter the floor-plate area and caused their premature turns along the ipsilateral floor-plate border (arrowhead in A) at 63.6±8.8% of the injection sites (n=7 embryos, 90 injection sites; G). These results are in line with previous reports (Stoeckli and Landmesser, 1995; Philipp et al., 2012). When both NrCAM and Endoglycan were downregulated, the number of ipsilateral turns was reduced to control levels (B,G; 7.3±5.8%, n=10 embryos; 78 injection sites), consistent with the idea that a decrease in adhesion due to a lack of NrCAM can be balanced by an increase in adhesion between floor plate and growth cones due to a lack of Endoglycan. Downregulation of either Podocalyxin (C) or CD34 (E) did not impair axon guidance (see also Figure 1; 12.3±5.2% (n=9) and 7.25±4.3% (n=8), respectively). In contrast to Endoglycan, neither concomitant downregulation of Podocalyxin (D) nor CD34 (F) could rescue the NrCAM-induced ipsilateral turns, as aberrant axon behavior was still observed at 56.7±9.2% (n=8 embryos, 77 injection sites) and 49.6±8.1% (n=10 embryos, 100 injection sites), respectively. The floor plate is indicated by dashed lines. For statistical analysis, one-way ANOVA followed by Tukey’s multiple comparisons test was used, ** p<0.01; (ns) p≥0.05. Bar: 50 µm.

### Endoglycan acts as ‘lubricant’ for growth cone movement in the floor plate

To get more insight into the role of Endoglycan in the regulation of contacts between axons and floor-plate cells, we established ex vivo live imaging of commissural axons during midline crossing. Intact spinal cords of HH22 chicken embryos, which were co-injected and unilaterally electroporated with constructs expressing farnesylated td-Tomato (td-Tomato-f) under the control of the dI1 neuron-specific Math1 enhancer together with farnesylated EGFP (EGFP-f) under the control of the β-actin promoter, were cultured and imaged for 24 hours (Figure 6). This method allowed us to follow the behavior and trajectories of the very first wave of single dI1 axons entering, crossing, and exiting the floor plate in a conserved environment (arrowheads, Figure 6A_1_-A_3_). The expression of EGFP-f in all transfected cells and brightfield images helped us to define the floor-plate boundaries (white dashed lines) and midline (yellow dashed lines in Figure 6B,C).

**Figure 6.**
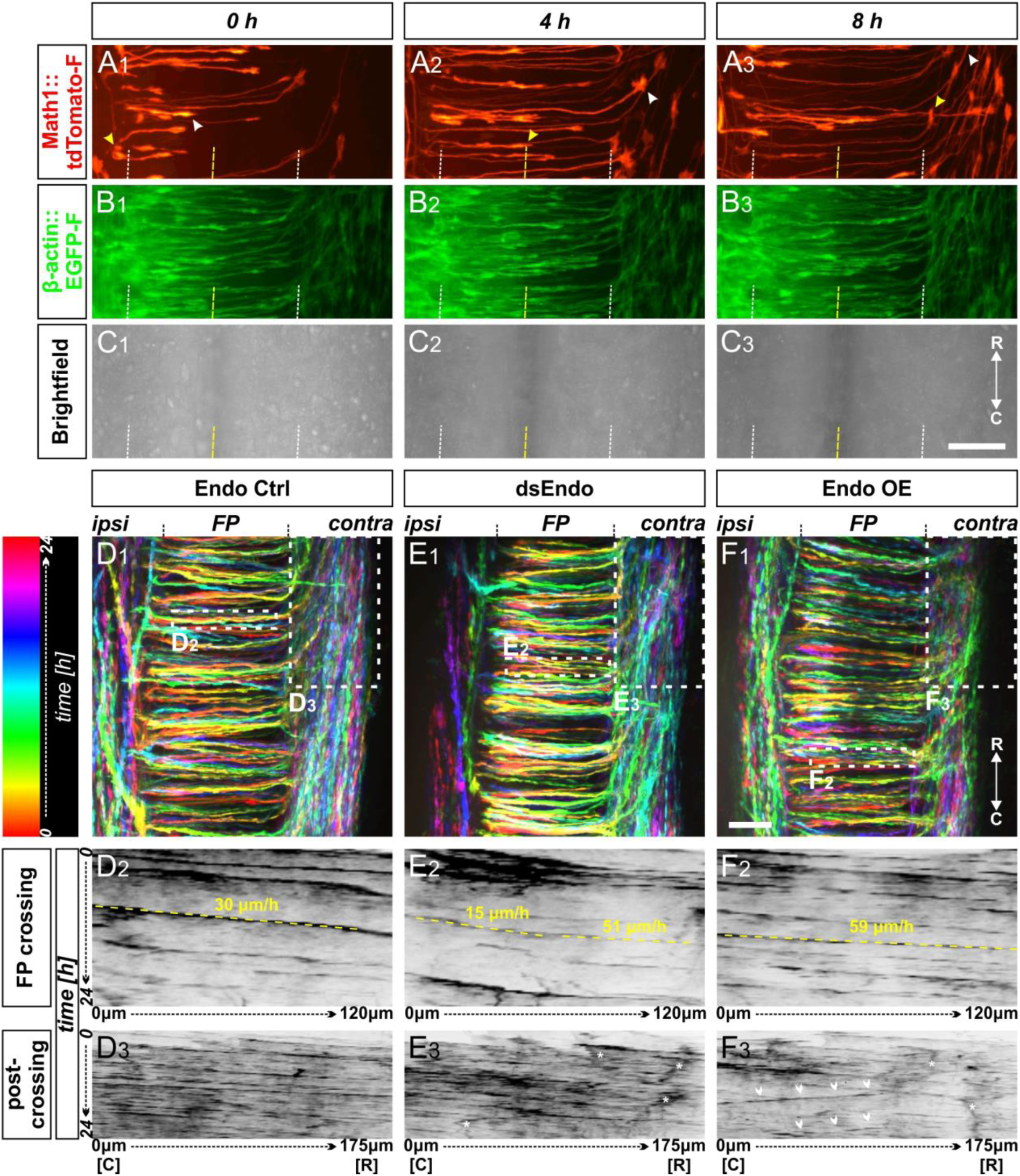
Live imaging of cultured intact spinal cords revealed major impacts of different Endoglycan levels on midline crossing. (A-C) Live imaging allowed tracing and quantitative analysis of dI1 axons’ trajectories in cultured intact chicken spinal cords. (A_1-3_) The behavior and trajectory of single tdTomato-positive dI1 axons could be tracked over time when they crossed the floor plate and turned rostrally (yellow and white arrowheads). (B-C) EGFP-F expression under the β-actin promoter and brightfield images helped to visualize the floor plate boundaries (white dashed line) and the midline (yellow dashed line). (D_1_, E_1_, F_1_) Temporally color-coded projections of 24h time-lapse movies (Supplementary movie 1). Kymograph analysis of the regions of interest selected in the floor plate of each condition shown in (D_1_,E_1_,F_1_) was used to calculate growth cone speed during floor-plate crossing (D_2_, E_2_, F_2_) and after turning into the longitudinal axis (D_3_, E_3_, F_3_). Yellow dashed lines outline a representative example of the slope (velocity) of a single axon crossing the floor plate in each condition. TdTomato-positive axons in control-injected spinal cords (D_1-3_) crossed the floor plate at a steady speed of 30 µm/h (D_2_) and turned rostrally in a highly organized manner (D_3_). In contrast, growth cone speed in the first half of the floor plate that was electroporated with dsRNA derived from Endoglycan (dsEndo) was markedly slowed down to only 15 µm/h. In the second, non-electroporated half of the floor plate, axons electroporated with dsEndo were faster than control axons (51 µm/h; E_2_). Axons overexpressing Endoglycan were faster in both halves of the floor plate (59 µm/h; F_2_). Downregulation or overexpression of Endoglycan clearly impacted the rostral turning behavior visualized by less organized patterns (D_3_-F_3_). Asterisks mark axons stalling and thus causing a ‘smeared’ pattern in the kymographs. Arrowheads indicate caudally turning axons. R, rostral; C, caudal; ipsi, ipsilateral; contra, contralateral; FP, floor plate. Scale bars: 50 µm.

Spinal cords of embryos co-injected with the Math1::tdTomato-f plasmid and dsRNA derived from Endoglycan (dsEndo) or a plasmid encoding chicken Endoglycan under the β-actin promoter (Endo OE) were imaged for 24 hours and compared to spinal cords dissected from control-injected embryos (Endo ctrl, Supplementary movie 1, temporally color-coded projections in Figure 6D_1_,E_1_ and F_1_). In contrast to control-injected spinal cords, the post-crossing segment of dI1 axons was disorganized in dsEndo and Endo OE conditions. In both these conditions, caudal turns were seen (Supplementary movie 1). As our in vivo data suggested a difference in adhesion between floor-plate cells and dI1 axons, we analyzed axonal midline crossing with kymographs in two different regions of interest (ROI; shown in Figures 6D_1_, E_1_, F_1_). This allowed us to follow growth cone movement across the floor plate and along the floor-plate border. Interestingly, our analyses indicated that the transfection of dsRNA derived from Endoglycan in dI1 neurons and floor-plate cells led to a decrease in the growth cones’ speed in the first half of the floor plate (15 µm/h) and in an increase in the second half (51 µm/h; Figure 6E_2_) compared to control-injected spinal cords (Figure 6D_2_ and Supplementary movie 2), where speed in the first and second halves did not differ (30 µm/h). In spinal cords overexpressing *Endoglycan*, growth cone speed was accelerated in the entire floor plate (59 µm/h; Figure 6F_2_; Supplementary movie 2). The analysis of axon growth in a second ROI confirmed the disorganization seen in both mutants in Supplementary movie 1. Whereas the axonal trajectories in control-injected embryos (Figure 6D_3_) were well organized and mostly parallel, axonal behavior in mutants caused ‘smeared’ patterns (asterisks Figure 6E_3_) due to stalling and pixels moving obviously in caudal direction indicating caudal axonal turns (arrowheads in Figures 6F_3_). These phenotypes confirmed our analyses of open-book preparations of spinal cords lacking Endoglycan (Figure 1). Axonal stalling (arrowheads) and caudal turns (arrows) could also be observed at the floor-plate exit site of spinal cord overexpressing *Endoglycan* (Supplementary movie 3).

The obvious differences in axonal behavior in experimental compared to control spinal cords was corroborated by quantitative analyses of specific aspects. Firstly, we quantified how much time the growth cones spent migrating from the floor-plate entry site to the exit site (Figure 7A_1_). Confirming the observations made in our movies and kymographic analysis, growth cones overexpressing *Endoglycan* crossed the floor plate faster, in only 4.4±1.4 hours (mean±SD), compared to controls (5.4±1.3 hours) and the dsEndo condition (5.5±1.2; Figure 7B). Furthermore, we compared the average time for crossing each half of the floor plate for each condition (Figure 7A_1_, C). Growth cones migrated equally fast through both halves in controls (2.7±0.9 hours versus 2.6±1.0 hours, Figure 7C). After overexpression of *Endoglycan*, there was a slight but significant shortening of the time growth cones spent crossing the first half compared to the second half (2.1±0.7 hours versus 2.3±0.8 hours, Figure 7C). In contrast, the unilateral silencing of *Endoglycan* induced a highly significant difference in the migration speed of growth cones in the first (electroporated) versus the second half of the floor plate. It took 3.1±0.9 hours to cross the first half but only 2.5±1.0 hours to cross the second half (Figure 7C). An alternative way of demonstrating the differences in migration speed is shown in Figure 7D,E. We calculated the ratios of the time spent in the first (Figure 7D) or the second half of the floor plate (Figure 7E) compared to the total time used for floor-plate crossing for the different conditions (Figure 7B). Indeed, knockdown of Endoglycan induced a significant increase in the ratio spent in the first half of the floor plate (0.56±0.1) compared to both control (0.51±0.1) and overexpression of Endoglycan (0.48±0.1; Figure 7D). In the second half of the floor plate, there was a significant decrease in spinal cords electroporated with dsEndo (0.44±0.1) compared to control (0.49±0.1) and *Endoglycan* overexpressing spinal cords (0.52±0.1; Figure 7E).

**Figure 7.**
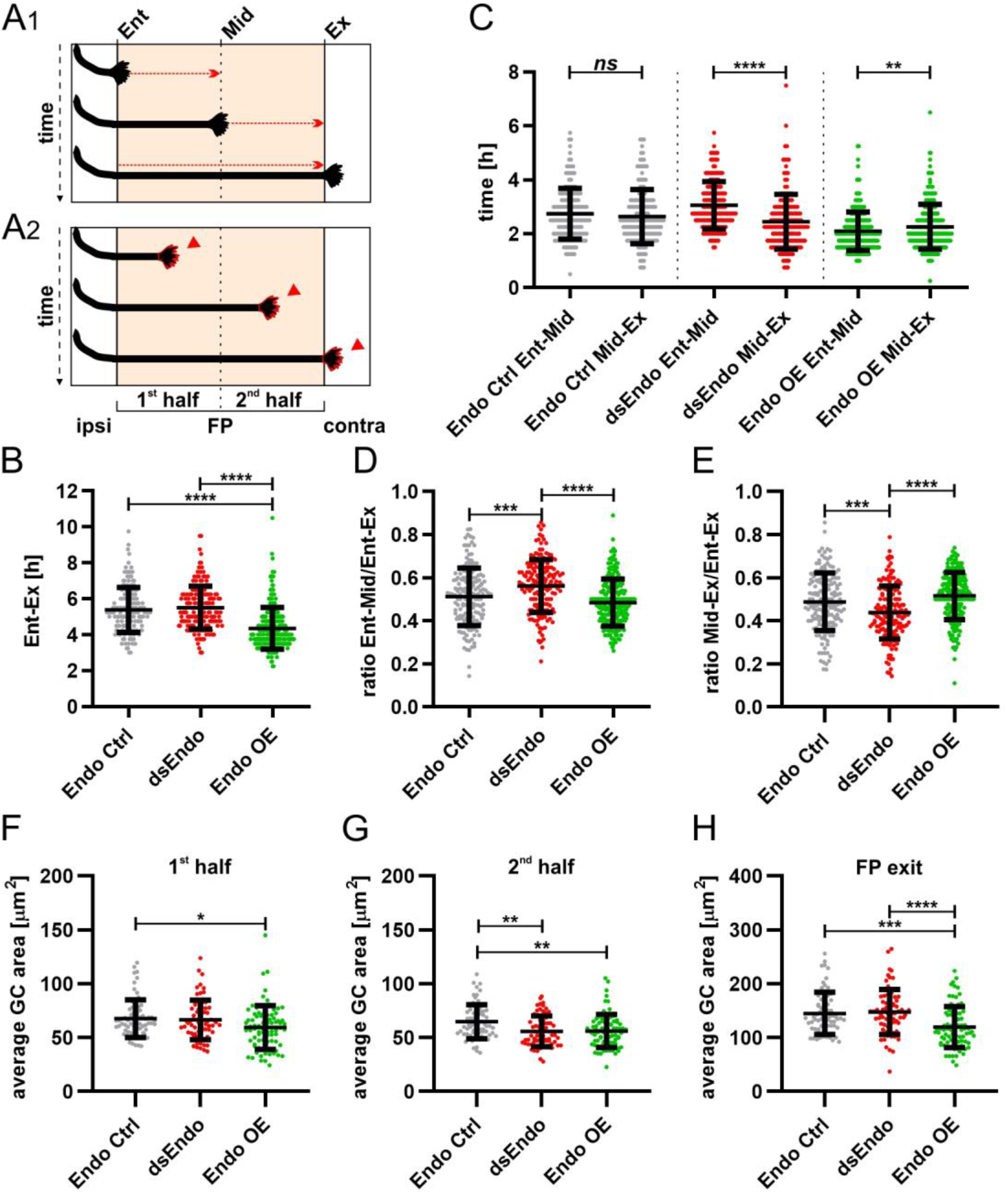
Too much or too little of Endoglycan impaired the timing and morphology of single dI1 growth cones migrating in the floor plate. Data at the single axon level extracted from 24h time-lapse recordings of tdTomato-positive dI1 axons crossing the floor plate. (A_1_) The time of floor-plate crossing was measured for the entire floor plate, for the first and for the second half for each condition. (A_2_) The average growth cone area was measured in the first half, the second half and at the exit site of the floor plate for each condition. (B) Overexpression of *Endoglycan* significantly decreased the time axons needed to cross the entire floor plate compared to control and dsEndo conditions (Kruskal-Wallis test with Dunn’s multiple-comparisons test). (C) The average time of crossing the first half and the second half of the floor plate was compared. Interestingly, there was a highly significant difference in the spinal cords unilaterally electroporated with dsEndo, as axons spent much longer in the first compared to the second half of the floor plate. There was no difference between the two halves of the floor plate in the control condition, but there was a significant decrease in growth cone speed between the first (electroporated) half of the floor plate and the second half, where only axons were overexpressing *Endoglycan* (Wilcoxon test). (D) and (E) The ratios of the time axons spent in the first half (D) or the second half (E) of the floor plate divided by the time they needed to cross it entirely were compared between conditions. Unilateral knockdown of Endoglycan resulted in a significant increase of the ratio in the first half and a decrease in the second half compared to both control and overexpression conditions (one-way ANOVA with Sidak’s multiple-comparisons test). (F-H) The average dI1 growth cone area at each position of the floor plate (as depicted in A_2_) were compared across all conditions. (F) Overexpression of Endoglycan induced a significant reduction in the average growth cone area compared to the control condition (Kruskal-Wallis test with Dunn’s multiple-comparisons test) but not compared to Endoglycan knockdown (*p* value = 0.08). (G) In the second half of the floor plate the average growth cone area was reduced in both Endoglycan knockdown and overexpression compared to control (one-way ANOVA with Sidak’s multiple-comparisons test). (H) At the floor plate exit site, overexpression of *Endoglycan* induced a significant decrease in the average growth cone area compared to both control and knockdown conditions (one-way ANOVA with Sidak’s multiple-comparisons test). Error bars represent standard deviation. p<0.0001 (****), p<0.001 (***), p<0.01 (**), p<0.05 (*) and p≥0.05 (ns) for all tests. See Table 2 for detailed results. Ent, entry; Mid, midline; Ex, exit; ipsi, ipsilateral; contra, contralateral; FP, floor plate; GC, growth cone.

Secondly, we also analyzed growth cone morphologies by comparing the average area in the first and the second halves, as well as at the exit site of the floor plate (Figure 7A_2_). The difference in growth speed was reflected in growth cone morphology and size (Supplementary Figure 8 and Supplementary movie 1). Growth cones tended to be small and have a simple morphology at fast speed. At choice points, like the floor-plate exit site, growth cone size and complexity increased. The average area of growth cones in the floor plate in control spinal cords was 68±18 µm^2^ in the first, and 65±16 µm^2^ in the second half of the floor plate (Supplementary Figure 8A). The growth cone area significantly increased at the floor-plate exit site, where growth cones need to choose to grow rostrally rather than caudally (Supplementary Figure 8A). At the exit site, growth cone area was 145±39 µm^2^ in control embryos. In agreement with their faster speed in the floor plate overexpressing *Endoglycan*, growth cones were significantly smaller in the first (60±20 µm^2^) and the second half (56±14 µm^2^) of the floor plate, as well as at the exit site (120±38 µm^2^) compared to controls (Figure 7F-H and Supplementary Figure 8C). Reduction of Endoglycan expression in the axons and in the first half of the floor plate resulted in reduced migration speed, but the average size of the growth cones was not significantly different from controls. (Figure 7F). The fact that growth cone were significantly faster in the second, non-transfected half of the floor plate was reflected by a significant reduction in growth cone area (56±14 µm^2^) compared to control (68±18 µm^2^) and compared to the first, transfected half (67±18 µm^2^; Figure 7G, Supplementary Figure 8B, Table 2).

**Table 2.**
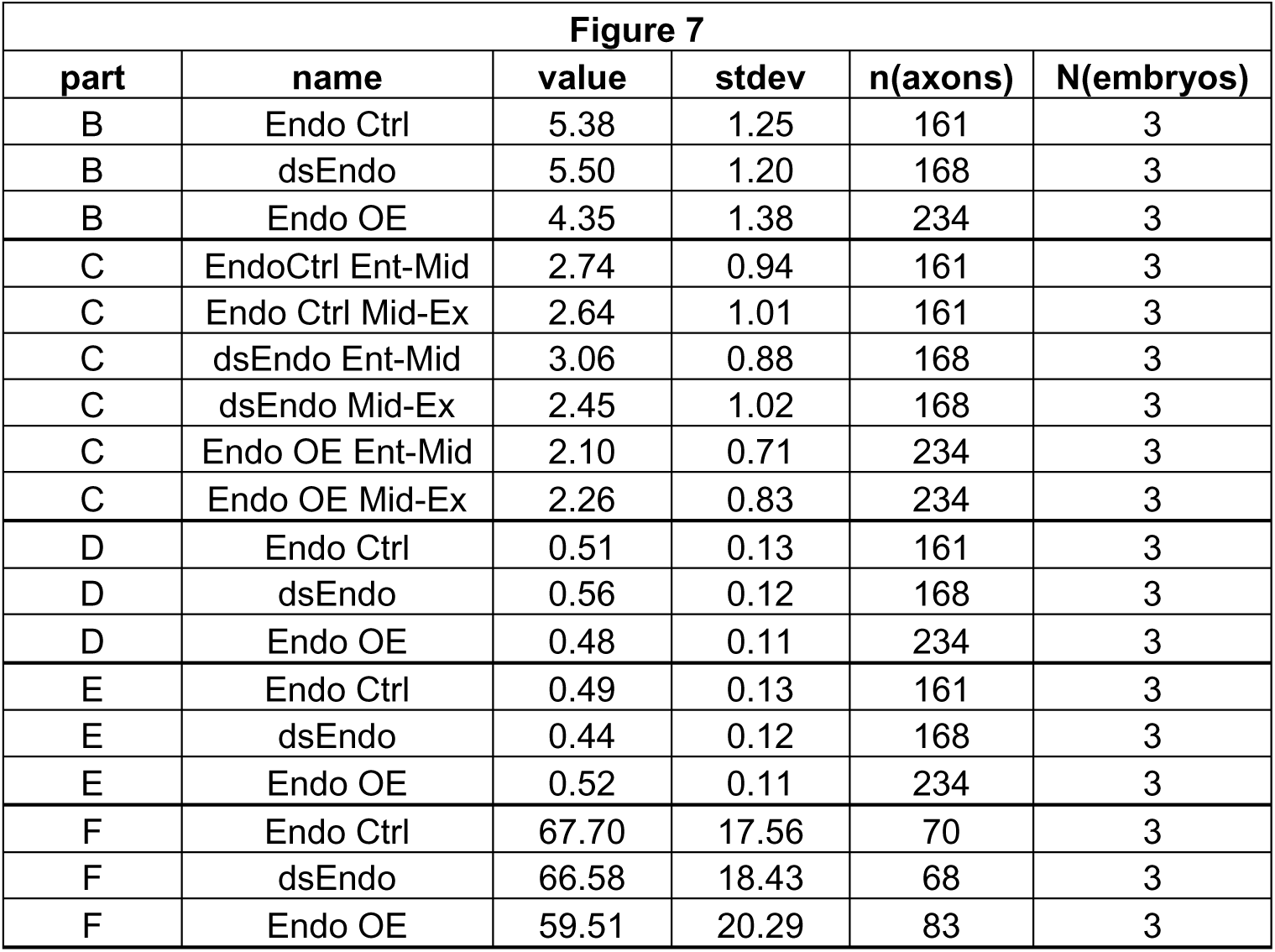
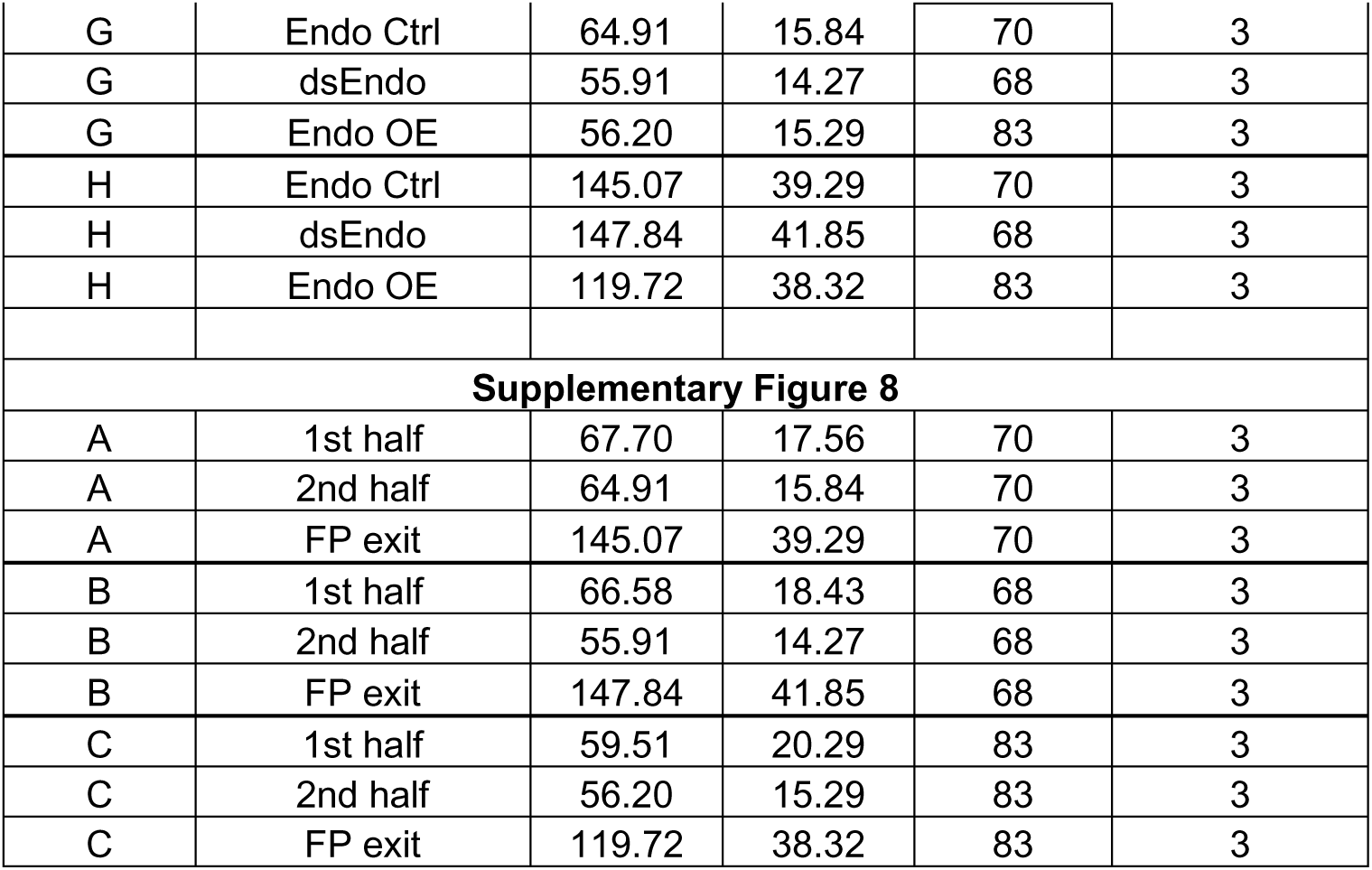
Quantification of midline crossing of dI1 axons using live imaging. The table contains the detailed values from results presented in Figure 7 and Supplementary Figure 8. Ent, entry; Mid, midline; Ex, exit; FP, floor plate; stdev, standard deviation.

Finally, live imaging of growth cones crossing the floor plate provided support for our hypothesis that axon-floor plate contact was causing the displacement of floor-plate cells observed after knockdown of *Endoglycan*, the ‘corkscrew’ phenotype. The tortuous, corkscrew-like phenotype of axons was seen exclusively in spinal cords after knockdown of *Endoglycan* (Figure 8 and Supplementary movies 4 and 5). We could observe roundish EGFP-f-positive cells (arrows) that obstructed the smooth trajectory of dI1 axons in the commissure (arrowheads in Figure 8A_1-5_ and Supplementary movie 4). Although we could not use markers to identify these cells as floor-plate cells, their position indicated that they had to be mislocalized floor-plate cells. Moreover, dI1 axons were found to form loops in the layer where floor-plate cell somata were localized (arrowhead in Figure 8B_1-5_ and Supplementary movie 5). In the first half of the floor plate electroporated with dsEndo, clusters of roundish EGFP-f-positive cells in the commissure (arrows) were apparently causing axons to deviate from their trajectory by strongly adhering to them (arrowheads in Figure 8C_1-5_ and Supplementary movie 5). We never found such aberrant behavior in control embryos or in embryos overexpressing *Endoglycan*. Moreover, we only observed these events after many dI1 axons had already crossed the floor plate (after at least 10 hours), supporting the hypothesis that the phenotype was due to excessive growth cone-floor plate adhesion resulting in floor-plate cell displacement. Furthermore, these observations suggest that Endoglycan regulates migratory speed of growth cones by modulating their adhesion to floor-plate cells.

**Figure 8.**
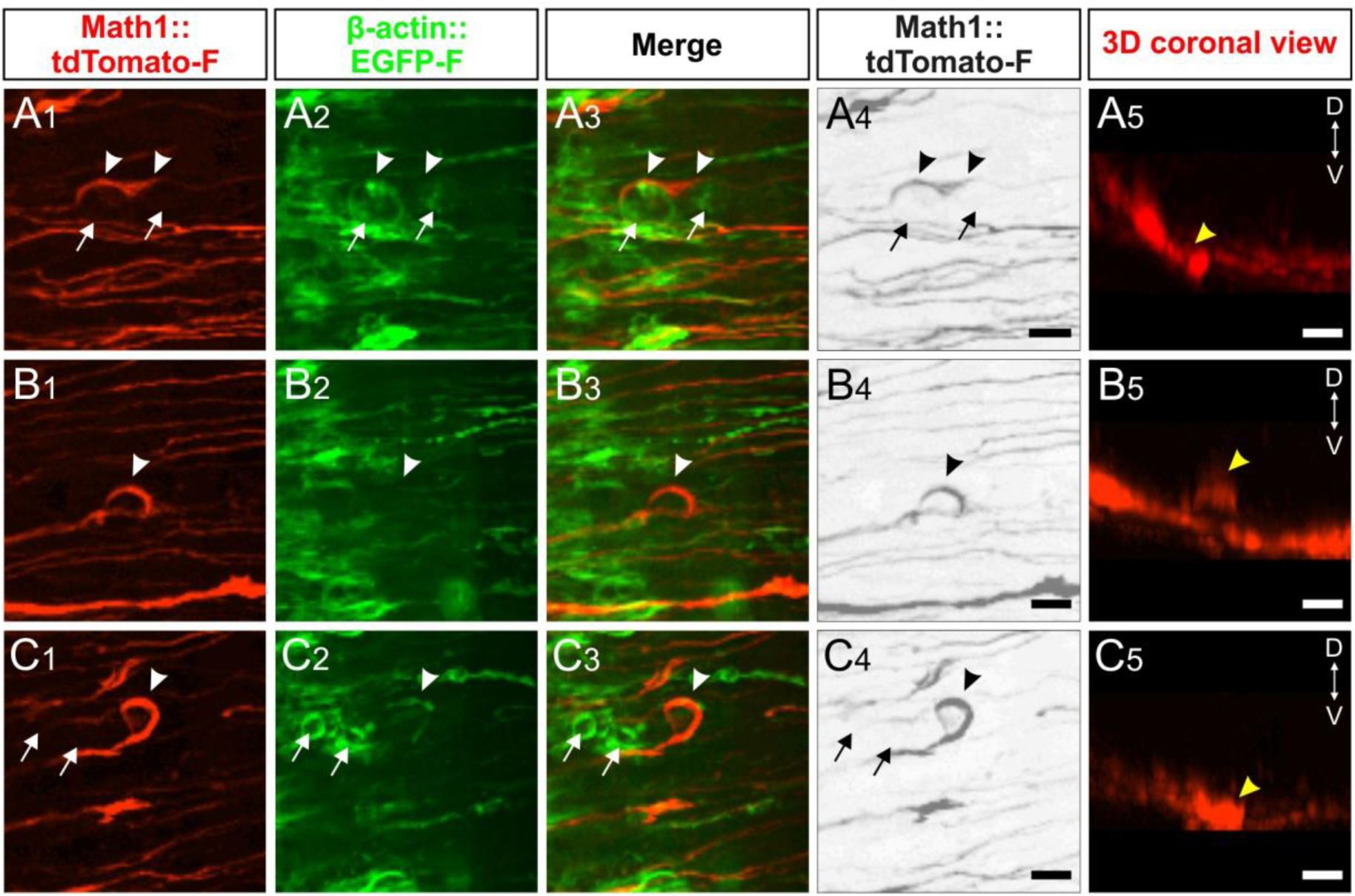
Live imaging of dI1 axons after perturbation of Endoglycan expression explains ‘corkscew-like’ phenotypes by aberrant interactions between axons and floor-plate cells. (A-C) ‘Corkscrew’-like phenotypes of dI1 axons expressing farnesylated tdTomato were observed by live imaging in the first half of the floor plate (electroporated half) only after *Endoglycan* was silenced (see also Supplementary movies 4 and 5). (A) Dislocated EGFP-positive cells (arrows) caused dI1 axons to deviate from a smooth trajectory inducing a ‘corkscrew-like’ morphology (arrowheads, A_1-4_). These axons and cells were located in the commissure as shown in a coronal view (yellow arrowhead, A_5_ and Supplementary movie 4). (B) Some axons were found to form a loop (arrowheads, B_1-4_) invading the layer of the floor plate where somata of floor-plate cells are located as shown in the transverse view (yellow arrowheads, B_5_ and Supplementary movie 5). (C) Clusters of dislocated roundish EGFP-positive cells (arrows) seemed to retain growth cones in the first half of the floor plate (arrowheads). These axons and cells were located in the commissure as shown in a coronal view (yellow arrowhead, C_5_ and Supplementary movie 5). D, dorsal; V, ventral. Scale bars: 10 µm.

Taken together, our live imaging studies support results from in vitro adhesion experiments indicating that the level of Endoglycan expression modulates adhesive strength between dI1 commissural growth cones and floor-plate cells. In contrast, axon-axon interactions did not seem to be different in the presence and absence of Endoglycan, as we did not find any effect on pre-crossing axons after perturbation of Endoglycan levels (Supplementary Figure 9).

### *Endoglycan* is expressed in migrating Purkinje cells and is required for their radial migration

In the developing cerebellum, *Endoglycan* expression was found exclusively in migrating Purkinje cells (Figure 9). Purkinje cells are born in the ventricular zone of the cerebellar anlage (Hatten, 1999). From there, they migrate radially toward the periphery to form the distinct Purkinje cell layer (Figure 9A-F). At HH44, when Purkinje cells migration is completed, *Endoglycan* mRNA is no longer detectable in Purkinje cells (Figure 9F). To test our hypothesis that Endoglycan was required as a ‘lubricant’ or regulator of cell-cell adhesion in a different situation, we analyzed the migratory behavior of Purkinje cells. To this end, we injected and electroporated dsRNA derived from *Endoglycan* into the developing cerebellum at HH34. Control embryos were injected and transfected only with a plasmid encoding EGFP (Figure 10). In untreated control embryos at HH38, the Purkinje cell layer is still more than one cell diameter in width but is clearly detectable in the periphery of the cerebellar lobes (Figure 10B,C). Very few, if any, Purkinje cells were found in the center of the lobes. The same was true in embryos injected with the EGFP-expression plasmid only (Figure 10D-F). In contrast, Purkinje cells were still found in the center of the lobes and close to the ventricular zone in HH38 embryos treated with dsRNA derived from *Endoglycan* (Figure 10G-I,J,M). In addition, the gross morphology of the cerebellum was severely compromised, because individual lobes failed to separate. Overall the size of the cerebellum was significantly reduced (Figure 10K,L,N,O).

**Figure 9.**
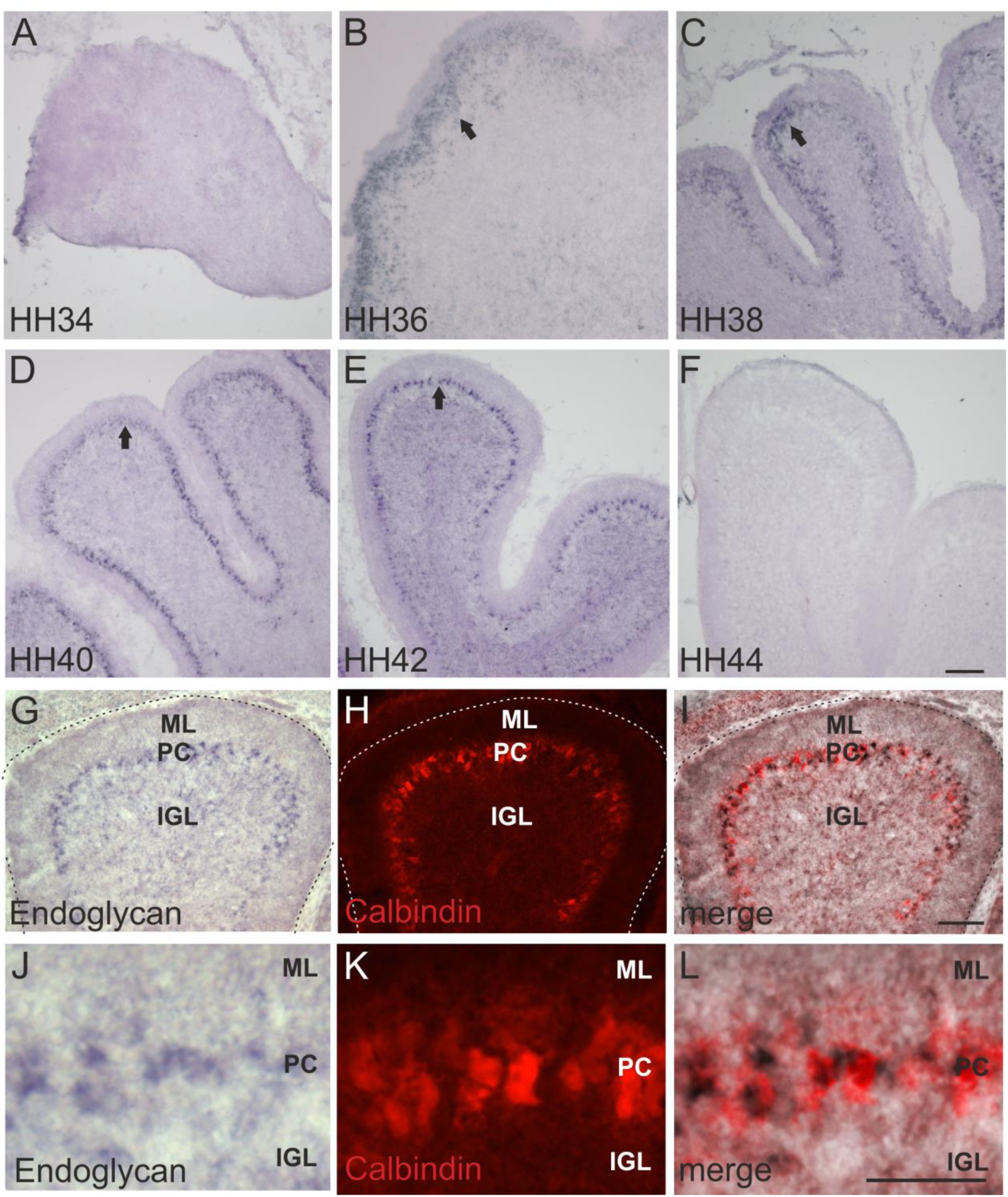
*Endoglycan* expression is restricted to Purkinje cells in the developing cerebellum. The temporal analysis of *Endoglycan* expression in the developing cerebellum localized it to migrating Purkinje cells. No *Endoglycan* expression was found at HH34 (A). Starting at HH36 (B), *Endoglycan* mRNA was found in migrating Purkinje cells. No change in expression was seen at HH38 (C), HH40 (D), and at HH42 (E) (arrows). At HH44, after migration of Purkinje cells was completed, *Endoglycan* was no longer expressed (F). We used Calbindin to identify Purkinje cells (H,K). At HH38, the in situ signal for *Endoglycan* (G,J) co-localized with Calbindin staining (H,K; overlay in I,L). Bar: 100 µm in A-F, 100 µm in G-I, 50 µm in J-L.

**Figure 10.**
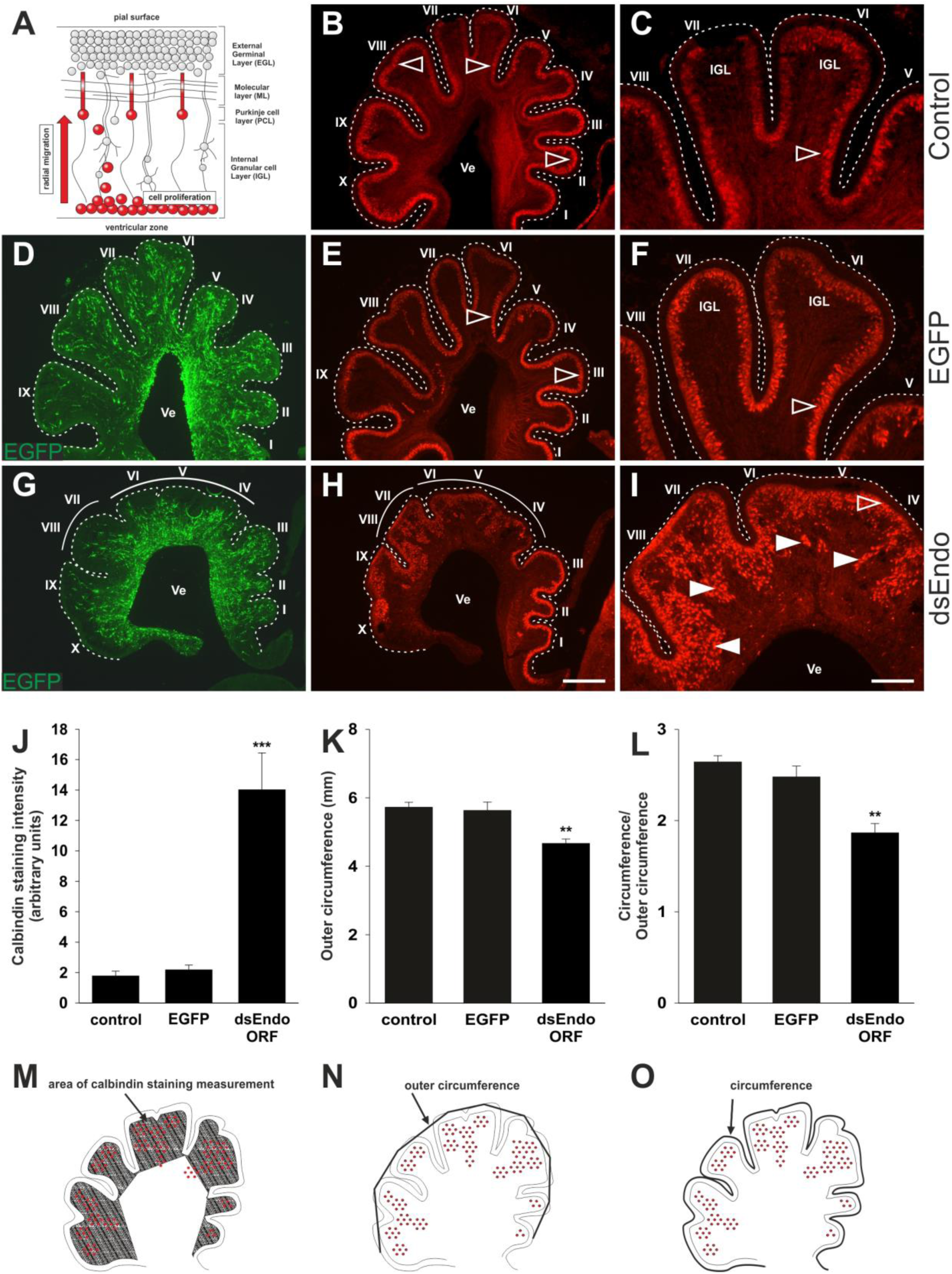
Endoglycan is required for Purkinje cell migration. Purkinje cells are born in the ventricular zone of the cerebellum. They migrate radially toward the periphery of the cerebellar folds to form the Purkinje cell layer (A). In control embryos at HH38, the Purkinje cell layer visualized by Calbindin staining is clearly detectable, although not fully matured to a monolayer (B and C). Control-injected embryos (D-F) were not different from untreated control embryos. Calbindin stains Purkinje cells in the periphery of the cerebellar folds (E and F). In the absence of Endoglycan (G-I) Purkinje cells failed to migrate and remained stuck in the center of the cerebellar folds (H, arrowheads in I). The failure of Purkinje cells to migrate radially was quantified by measuring fluorescence intensity for Calbindin in control (n=5 embryos), *EGFP*-injected (n=4 embryos), and *Endoglycan* dsRNA-treated embryos (J; n=5 embryos). Calbindin staining intensity was measured as indicated in (M). The increase is highly significant for *Endoglycan* dsRNA-treated embryos, p < 0.001. As a measure for the size of the cerebellum, the outer circumference was measured as indicated in (N). The cerebellum was smaller in experimental embryos compared to both control groups (K; p<0.01). In order to quantify the reduction in the number of folds that was obvious from the visual inspection of cerebellar sections, we measured the actual circumference of the cerebellum as indicated in (O) and divided it by the outer circumference. Control embryos had a ratio of 2.64±0.06 and 2.47±0.11, respectively. Embryos lacking Endoglycan showed a clear reduction in the ratio between the two circumferences with a value of 1.8±0.1 (L; p<0.01), indicating that they had fewer cerebellar folds compared to control embryos, which had always 10 distinct folds. Bar: 500 µm in B, D, E, G, H; 200 µm in C, F, I. ANOVA with Bonferroni post-hoc test was used for statistical analysis.

### Aberrant migration of Purkinje cells reduces granule cell proliferation

Purkinje cells regulate the proliferation of granule cells by releasing Shh (Sonic hedgehog), which stimulates proliferation of granule cell precursors in the outer EGL (external granule cell layer) (Dahmane and Ruiz i Altaba, 1999; Wechsler-Reya and Scott, 1999; Wallace, 1999; Lewis et al., 2004; reviewed in De Luca et al., 2016). In turn, reduced proliferation of granule cells was shown to result in changes of cerebellar morphology similar to the ones we observed after downregulation of Endoglycan (Figure 10) (Lewis et al., 2004). A reduced rate of granule cell proliferation was indeed what we found in embryos after silencing *Endoglycan*. When we used Pax6 as a marker for granule cells, we found a thinner EGL in experimental embryos compared to control-treated and untreated embryos (Figure 11A-D). This decrease in EGL width was due to a reduced proliferation rate of granule cells rather than apoptosis (Figure 11E-H). In contrast to granule cells, the proliferation rate of Purkinje cells and other cells born in the ventricular zone at HH35 did not differ between control embryos and embryos lacking Endoglycan (Figure 11I-K).

**Figure 11.**
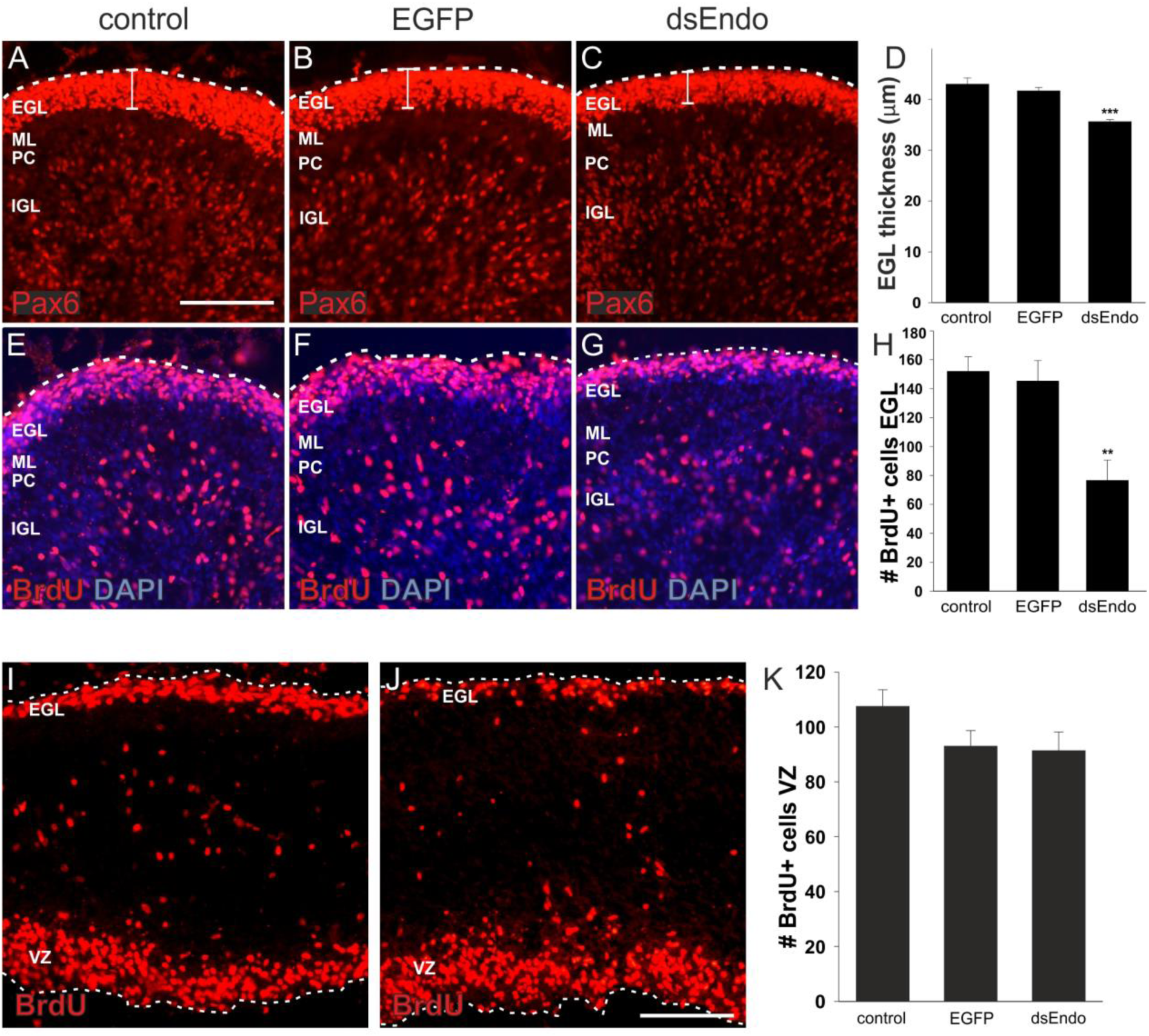
Downregulation of Endoglycan results in a smaller cerebellum due to reduced proliferation of granule cells. The failure of Purkinje cell migration has negative consequences on granule cell proliferation (A-H). Granule cells in the EGL and in the developing IGL are labeled by Pax6 (A-C). A reduction in the width of the EGL was found for embryos treated with dsRNA derived from *Endoglycan* (C,D; n=3 embryos; p<0.001) compared to age-matched untreated embryos (A; n=4 embryos), or control-treated embryos, expressing EGFP (B; n=3 embryos), when sections from the same relative position of the cerebellum were analyzed. No difference was found between the two control groups (D). The proliferation of granule cells in the outer EGL was visualized by BrdU incorporation (E-H). Embryos were exposed to BrdU for 3 hours before they were sacrificed at HH38. The number of BrdU-positive cells in the outer EGL was compared between untreated (E; n=6 embryos), EGFP-expressing control embryos (F; n=4 embryos) and embryos lacking Endoglycan (G; n=5 embryos). The number of BrdU-positive cells was significantly reduced in embryos lacking Endoglycan (H; p<0.01). The number of BrdU-positive cells in the ventricular zone at HH35 did not differ between untreated controls (I) and embryos lacking Endoglycan (J; K). The number of BrdU-positive cells in the ventricular zone is given per 10’000 µm^2^. Bar: 100 µm. ANOVA with Bonferroni post-hoc test was used for statistical analysis.

In summary, our results demonstrate a vital role for Endoglycan in commissural axon guidance at the ventral midline and Purkinje cell migration in the developing cerebellum. In both systems, the observed phenotype is consistent with the hypothesis that Endoglycan is an essential regulator of cell-cell contacts by modulating the strength of adhesion between cells. This model is supported by observations in vitro and in vivo. Neuronal attachment was negatively affected by the presence of an excess of Endoglycan in a glycosylation-dependent manner, indicating that Endoglycan decreases adhesive strength during neural circuit assembly.

## DISCUSSION

We identified *Endoglycan*, a member of the CD34 family of sialomucins, in a screen for axon guidance cues involved in commissural axon pathfinding at the midline of the spinal cord. In the developing chicken cerebellum, *Endoglycan* is expressed exclusively in Purkinje cells during their migration from the ventricular zone to their final destination in the periphery of the cerebellar lobes (Figure 9). In the absence of Endoglycan, Purkinje cells failed to migrate and accumulated in the center of the cerebellar folds (Figure 10). This observation suggests a role of Endoglycan as ‘lubricant’ that is supported by the structural features of sialomucins. The function of CD34 family members has not been characterized in detail but all the results obtained so far are compatible with an anti-adhesive role (Nielsen and McNagny, 2008). One exception are reports from the lymph node cells, the so-called high endothelial venules (HEVs) where a very specific glycosylation patterns was implicated in the interaction of CD34 and Endoglycan with L-selectin (Furness and McNagny, 2006). However, in agreement with most published studies on the role of CD34 and Podocalyxin (for reviews see Furness and McNagny, 2006; Nielsen and McNagny, 2008 and 2009) our observations suggest that Endoglycan acts as a ‘lubricant’ rather than as adhesive factor. This model is supported by results from in vivo and in vitro experiments that confirm a negative effect of Endoglycan on cell-cell adhesion (Figure 12).

**Figure 12.**
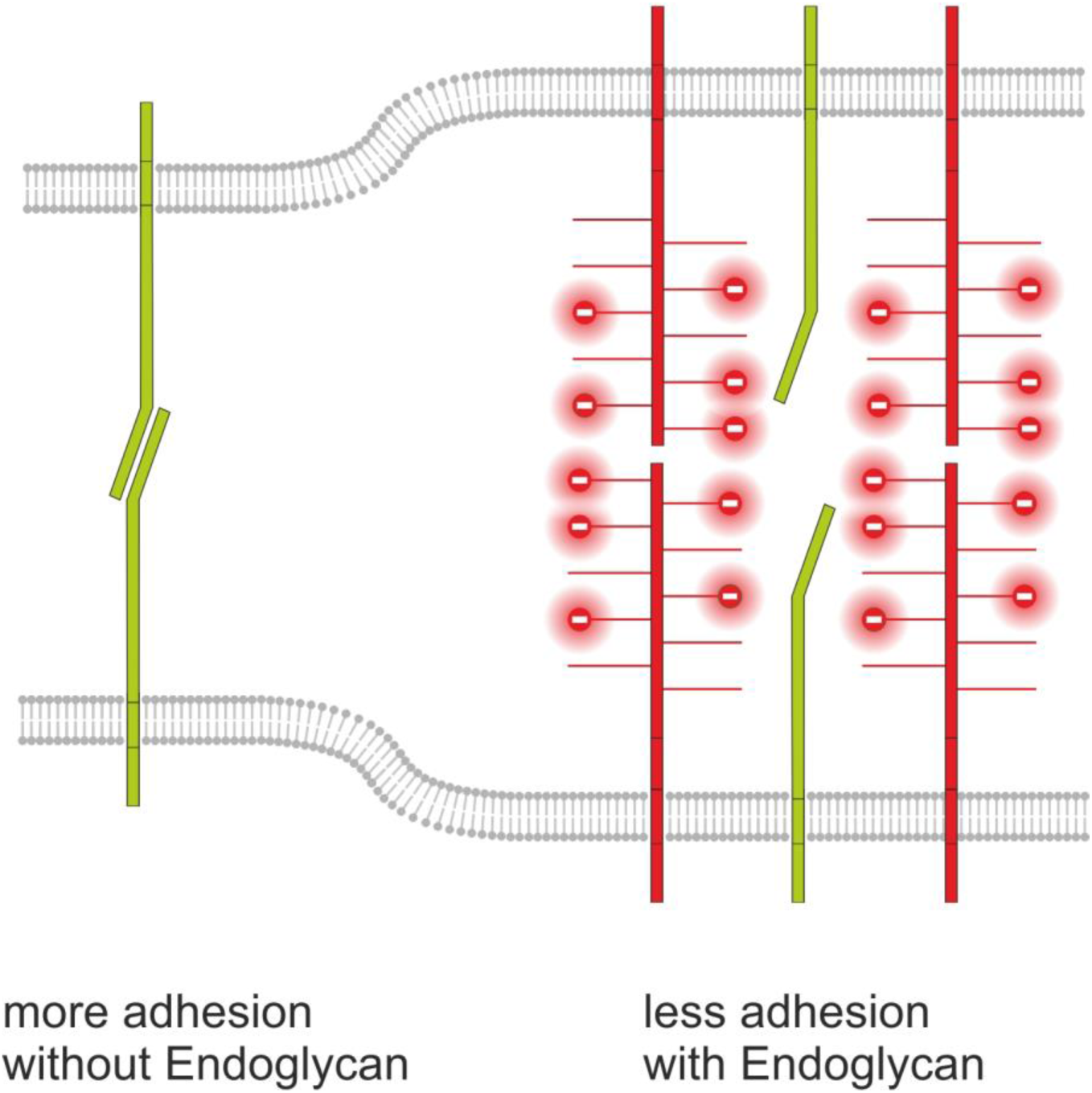
Endoglycan modulates cell-cell contact by interference with adhesive strength. Based on our in vivo and in vitro studies we postulate a model for Endoglycan function in neural circuit formation that suggests an ‘anti-adhesive’ role by modulation of many specific molecular interactions due to decreasing cell-cell contact. This model is consistent with our rescue experiments demonstrating that the source of Endoglycan did not matter but the expression level did, as aberrant phenotypes were prevented when Endoglycan was expressed either in the axon or in the floor plate.

The adhesion-modulating effect of Endoglycan is mediated by the negatively charged mucin domain. Similar to the role suggested for the polysialic acid modification of NCAM (Rutishauser, 2008; Brusés and Rutishauser, 2001; Burgess et al., 2008), Endoglycan could lower cell-cell adhesion by increasing the distance between adjacent cell membranes due to repulsion caused by the bulky, negatively charged posttranslational modifications of its extracellular domains. A similar effect was found for PSA-NCAM in hindlimb innervation (Tang et al., 1994; Landmesser et al., 1990) and in the visual system, where retinal ganglion cell axons innervating the tectum were found to regulate axon-axon adhesion versus axon-target cell adhesion (Rutishauser et al., 1988). The same mechanism was found in motoneurons, where axon-axon versus axon-muscle fiber adhesion was a determining factor for the appropriate innervation pattern. In contrast to PSA-NCAM that continues to play a role in synaptic plasticity in the adult nervous system, the function of Endoglycan appears to be restricted to development. Expression of *Endoglycan* ceased in the cerebellum after the mature wiring pattern was achieved.

At first sight, the effect of Endoglycan on floor-plate morphology appears to suggest a positive regulation of cell-cell adhesion. Floor-plate cells are precisely aligned in control embryos but are protruding into the commissure in the absence of Endoglycan. Therefore, one might conclude that in the absence of Endoglycan cell-cell adhesion between floor-plate cells is compromised, resulting in the observed structural changes. However, this scenario can be ruled out based on the analysis of younger embryos. At HH21, the floor plate was intact in the absence of Endoglycan, indicating that Endoglycan is not required for adhesion between floor-plate cells. The morphology of the floor plate is only compromised once many axons have crossed the midline. These findings are supported by our live imaging data of growth cones crossing the floor plate. Contacts between commissural axons and floor-plate cells have to be broken when later crossing commissural axons arrive and cross (Yaginuma et al., 1991). Commissural axons crossing the floor plate are suggested to do so by close interaction with short filopodial processes of floor-plate cells. Thus, the aberrant morphology of the floor plate at HH25 is explained by the inability of axons to break contacts with floor-plate cells in the absence of Endoglycan, consistent with our hypothesis that Endoglycan is a negative regulator of adhesion by acting as a ‘lubricant’. Live imaging data demonstrated that the perturbation of the balance in growth cone-floor plate adhesion led to impaired timing of midline crossing, which in turn might also interfere with the correct sensing and reading of guidance cues by dI1 growth cones, and prevented them from making the correct decision at the floor-plate exit site.

Thus, we concluded that the function of Endoglycan in commissural axon guidance and in Purkinje cell migration is to lower cell adhesion. In both cases, the absence of Endoglycan results in too much stickiness. In the cerebellum, excessive adhesion prevents the Purkinje cells from migrating to their target layer. At the midline of the spinal cord, excessive adhesion causes axons to adhere too much to floor-plate cells and prevents their displacement by follower axons. Rather than acting as a guidance cue or guidance receptor, we suggest that Endoglycan affects neural circuit formation by modulating the interaction of many different guidance cues and their surface receptors.

In summary, we propose an ‘anti-adhesive’ role for Endoglycan in axon guidance and neural migration that is fine-tuning the balance between adhesion and de-adhesion (Figure 12). Precise regulation of cell-cell contacts is required in both processes and is fundamental for developmental processes that depend on a high degree of plasticity and a plethora of specific molecular interactions.

## METHODS

### Identification and Cloning of *Endoglycan*

We had used a PCR-based subtractive hybridization screen to search for guidance cues for post-crossing commissural axons (for details, see Bourikas et al., 2005). To this end, we isolated floor-plate cells from HH26 and HH20 embryos (Hamburger and Hamilton, 1951). Among the differentially expressed genes, we found a sequence from the 3’-UTR of *Endoglycan*. Subsequently, a cDNA fragment from the coding sequence of *Endoglycan* (PODXL2; 1028-1546 bp) was obtained by RT-PCR using total RNA isolated from HH40 cerebellum. For reverse transcription, 1 µg total RNA was mixed with 0.3 µl RNasin (Promega), 1 µl dNTPs (5 mM), 1 µl random nonamers, 1 µl DTT (Promega), in 20 µl Superscript II buffer (Invitrogen). Reverse transcription was carried out for 1 hour at 42°C. Two µl of this mixture were used for PCR with 2.5 µl forward primer (10 µM; 5’-CAGACACGCAGACTCTTTC-3’) and 2.5 µl reverse primer (10 µM; 5’-CTAAAGATGTGTGTCTTCCTCA-3’) using the Expand Long Template PCR System (Roche). The PCR conditions were 35 cycles at 95°C for 30 sec, 57°C for 30 sec and 68°C for 3 min. The PCR product was cut with BamHI/BclI and cloned into pBluescript II KS. For cloning of full-length chicken Endoglycan, we used 5’-ATGGTGAGAGGAGCTGCG-3’ and 5’-GTGTTTGAGGAAGACACACATCTTTAG-3’ as forward and reverse primers, respectively. A plasmid containing the full-length ORF of human *Endoglycan* was obtained from SourceBiosource.

### Preparation of DIG-labeled RNA probes and in situ hybridization

For in vitro transcription, 1 µg of the linearized and purified plasmids encoding Endoglycan (EndoORF: 1028-1546pb, Endo3’UTR: 3150-3743bp and 5070-5754bp; numbers are derived from the human sequence), Podocalyxin (ChEST190L9) and CD34 (ChEST91D7) were used to prepare DIG-labeled in situ probes as described earlier (Mauti et al., 2006). The same fragments were used to prepare dsRNA (Pekarik et al., 2003; Baeriswyl et al., 2008; Andermatt and Stoeckli, 2014b).

### Northern Blot

Total RNA was extracted from cerebrum, cerebellum, spinal cord, muscle, heart, lung and kidney from HH38 embryos using the RNeasy Mini Kit (Qiagen) and loaded on a denaturing formaldehyde gel (4.5 µg of total RNA per lane). The RNA was blotted onto a positively charged nylon membrane (Roche) overnight using 10x SSC as a transfer medium. The membranes were hybridized with 1.5 µg preheated DIG-labeled RNA probes for *Endoglycan* and *GAPDH* at 68°C overnight. The membrane was then washed twice with 2xSSC/0.1%SDS for 5 minutes at room temperature and twice with 0.1xSSC/0.5% SDS for 20 minutes at 68°C. For detection, buffer 2 (2% blocking reagent dissolved in 0.1 M maleic acid, 0.15 M NaCl, pH 7.5) was added for 2-3 hours at room temperature. After incubation with anti-digoxigenin-AP antibody dissolved in buffer 2 (1:10,000; Roche) for 30 minutes at room temperature the membrane was washed twice in washing buffer (0.3% Tween 20 dissolved in 0.1 M maleic acid, 0.15 M NaCl, pH 7.5) for 20 minutes. Subsequently, detection buffer (0.1 M Tris-HCl, 0.1 M NaCl, pH 9.5) was applied for 2 minutes before adding CDP-star (25 mM, Roche) for 5 minutes in the dark. For detection of the chemiluminescence a Kodak BioMAX XAR film was used.

### In ovo RNAi

For functional studies in the spinal cord, we silenced *Endoglycan* with three different long dsRNAs. They were produced from bp 1028-1546 of the ORF, as well as bp 3150-3743 and bp 5070-5754 from the 3’UTR. The fact that we obtained the same phenotype with three different, non-overlapping dsRNAs derived from *Endoglycan* confirms the specificity of the approach and the absence of off-target effects. dsRNA was produced as detailed in Pekarik et al., 2003 and Wilson and Stoeckli, 2011. Because no antibodies recognizing chicken Endoglycan are available, we used in situ hybridization to assess the successful downregulation of the target mRNA (Supplementary Figure 10). Downregulation efficiency was about 40%. Because we transfect only around 50% of the cells in the electroporated area, transfected cells express only very low levels of Endoglycan.

For rescue experiments, the dsRNA was co-injected with 150 (low), 300 (middle), or 750 ng/µl (high) plasmid encoding the ORF of chicken *Endoglycan*. The ORF was either expressed under the control of the Math1 promoter for dI1 neuron-specific expression, or the Hoxa1 promoter for floor-plate specific expression of *Endoglycan*.

### Ex ovo RNAi

To analyze the in vivo function of Endoglycan in the developing cerebellum, ex ovo cultures of chicken embryos were prepared (Baeriswyl and Stoeckli, 2008; Andermatt and Stoeckli, 2014b). Injections and electroporations were performed at E8 (HH34). To have direct access to the embryo a small hole of 3 to 4 mm diameter was cut into the extraembryonic membranes above the eye. For positioning and stabilization of the head during injection and subsequent electroporation, we used a hook prepared from a spatula. Approximately 1 µl of the nucleic acid mixture, consisting of a plasmid encoding EGFP under the control of the β-actin promoter (100 ng/µl), dsRNA derived from the ORF of *Endoglycan* (500 ng/µl), and 0.04% (vol/vol) Trypan Blue (Invitrogen) dissolved in sterile PBS was injected into the cerebellum, using a borosilicate glass capillary with a tip diameter of 5 µm (World Precision Instruments). Before electroporation, a few drops of sterile PBS were added to the embryo. For the electroporation, a platelet electrode of 7 mm diameter (Tweezertrodes Model #520, BTX Instrument Division, Harvard Apparatus) was placed collaterally to the head of the embryo. Six pulses of 40 V and 99 ms duration were applied using a square wave electroporator (ECM830, BTX).

### Motoneuron adhesion assay

Dissociated motoneurons of HH26 chicken embryos were cultured as described previously (Mauti et al., 2006) either on HEK293T cells stably expressing human Endoglycan-myc under the control of the CMV promoter or on untransfected HEK293T cells as control. The plasmid encoding human Endoglycan was obtained from SourceBioScience (Nottingham, UK). After 40h, the cultures were fixed for 1h at room temperature in 4% paraformaldehyde and stained with mouse anti-neurofilament (RMO 270; Zymed) and rabbit anti-myc antibodies (Abcam). The number of neurofilament-positive cells was counted in 16 randomly selected frames (0.4 mm^2^). Similar results were obtained in 3 independent experiments. One representative example is shown in Figure 4. In a separate set of experiments, cultured HEK cells expressing Endoglycan were treated with O-glycosidase (8’000 U/ml; NEB) or α2-3,6,8 Neuraminidase (5 U/ml; NEB) for 2 hours before motoneurons were added (Supplementary Figure 7).

### Tissue preparation and analysis

To analyze commissural axon growth and guidance the embryos were sacrificed between HH25 and 26. The spinal cord was removed, opened at the roof plate (‘open-book’ preparation) and fixed in 4% paraformaldehyde (PFA) for 40 min to 1 hour at room temperature. To visualize the trajectories of commissural axons, Fast-Dil (5 mg/ml, dissolved in ethanol, Molecular Probes) was injected into the dorsal part of the spinal cord as described previously (Wilson and Stoeckli, 2012). For the analysis of the cerebellum, the embryos were sacrificed one to four days after electroporation. The whole brain was removed and analyzed for EGFP expression using a fluorescence stereomicroscope (Olympus SZX12). The brain tissue was fixed for two hours at room temperature in 4% PFA in PBS. After fixation, the brain tissue was rinsed in PBS and transferred to 25% sucrose in 0.1M sodium phosphate buffer, pH 7.4, for cryoprotection. In this study, 30 µm-thick sagittal cryostat sections were used for analysis. For the preparation of cryostat sections, the brains were embedded in O.C.T Tissue-Tek (Sakura) in Peel-a-Way^®^ disposable embedding molds (Polysciences), frozen in isopentane on dry ice and cut on a cryocut (CM1850, Leica Microsystems). The sections were collected on SuperFrost^®^Plus microscope slides (Menzel-Glaeser).

### Immunohistochemistry

Cryostat sections were rinsed in PBS at 37°C for 3 minutes followed by 3 minutes in cold water. Subsequently the sections were incubated in 20 mM lysine in 0.1 M sodium phosphate (pH 7.4) for 30 minutes at room temperature before being rinsed in PBS three times for 10 minutes. The tissue was permeabilized with 0.1% Triton in PBS for 30 minutes at room temperature and then washed again three times with PBS for 10 minutes. To prevent unspecific binding of the antibody the tissue was blocked with 10% fetal calf serum (FCS) in PBS for one hour. Rabbit anti-GFP (1:250; Abcam), anti-axonin-1 (rabbit 1:1000 or goat 1:500), anti-NrCAM (goat 1:1000), anti-Calbindin D-28K (1:2000, CB38a; Swant), mouse anti-HNF3β (supernatant; 4C7, DSHB), mouse anti-Islet1 (supernatant; 40.2D6, DSHB), mouse anti-Nkx2.2 (supernatant; 74.5A5, DSHB), mouse anti-Pax3 and Pax6 (supernatant, DSHB) were dissolved in 10% FCS/PBS and incubated overnight at 4°C. After three washes in PBS, 10% FCS in PBS was applied again for one hour, followed by the incubation with goat anti-rabbit IgG-Alexa488 (1:250; Molecular Probes), donkey anti-rabbit IgG-Cy3 (1:200; Jackson ImmunoResearch) or goat anti-mouse IgG-Cy3 (1:250; Jackson ImmunoResearch) diluted in 10% FCS in PBS for 90 minutes at room temperature. The tissue was rinsed 5 times in PBS for 12 minutes and then mounted in Celvol (Celanese) or Mowiol. The staining of cryostat sections was analyzed with an upright microscope equipped with fluorescence optics (Olympus BX51).

### Analysis of cell proliferation and cell death

To assess cell proliferation in the developing cerebellum, we used BrdU incorporation. Embryos were injected and electroporated at HH34 with dsRNA derived from *Endoglycan* and the *EGFP* plasmid or with the *EGFP* plasmid alone. After 1 (HH35) or 4 days (HH38), 200 µl 50 mM BrdU in H_2_O were pipetted onto the chorioallantois. After 3 h the embryos were sacrificed, the brains were dissected and prepared for cryostat sections as described above. For visualization of the incorporated BrdU, the sections were incubated in 50% formamide in 2xSSC for 1 to 2 h at 65 °C, rinsed twice in 2xSSC for 15 min followed by incubation in 2 N HCl for 30 min at 37 °C. Sections were rinsed in 0.1 M borate buffer (pH 8.5) for 10 min at room temperature, followed by PBS (six changes). BrdU was detected with mouse anti-BrdU (Sigma; 1:200) using the protocol detailed above. Sections were counterstained with DAPI (5 µg/ml in PBS) for 20 min at room temperature.

Apoptosis was analyzed as described previously (Baeriswyl and Stoeckli, 2008). For analysis of cell death in the floor plate, we used cleaved caspase-3 staining of sections taken from HH25 embryos.

### Quantification

Dil injections sites with pathfinding errors were analyzed and counted, using an upright microscope equipped with fluorescence optics (Olympus BX51). All measurements including floor-plate width, thickness of the commissure, spinal cord width, Calbindin fluorescence intensities, real and outer cerebellar circumference, EGL thickness, and number of BrdU positive cells were performed with the analySIS Five software from Soft Imaging System. For all measurements, embryos injected with dsRNA derived from *Endoglycan* were compared with embryos injected with the EGFP plasmid only, and untreated controls. For statistical analyses, ANOVA with Bonferroni correction was used except for the rescue experiments, where Tukey’s multiple comparison test was used instead. Values are given as mean ± SEM. *: P < 0.05. **: P < 0.01. ***: P < 0.001.

### Live imaging

Plasmids encoding farnesylated td-Tomato under the Math1 enhancer upstream of the β-globin minimal promoter for dI1 neuron-specific expression (Math1::tdTomato-f) and farnesylated EGFP under the β-actin promoter (β-actin::EGFP-f) were co-injected into the central canal of the chicken neural tube in ovo at HH17/18 and unilaterally electroporated, using a BTX ECM830 square-wave electroporator (five pulses at 25 V with 50 ms duration each), as previously described (Wilson and Stoeckli, 2012). For the perturbation of Endoglycan levels, either 300 ng/µl dsRNA derived from the 3’-UTR of Endoglycan or a plasmid encoding the open-reading frame of Endoglycan under the β-actin promoter were co-injected with the Math1::tdTomato-f plasmid. After electroporation, embryos were covered with sterile PBS and eggs were sealed with tape and incubated at 39°C for 26-30 hours until embryos reached stage HH22.

For live imaging, embryos were sacrificed at HH22. Intact spinal cords were dissected and embedded with the ventral side down in a drop (100 µl) of 0.5% low-melting agarose (FMC; Pignata et al., 2019) containing a 6:7 ratio of spinal cord medium (MEM with Glutamax (Gibco) supplemented with 4 mg/ml Albumax (Gibco), 1 mM pyruvate (Sigma), 100 Units/ml Penicillin and 100 µg/ml Streptomycin in a 35-mm Ibidi µ-Dish with glass bottom (Ibidi, #81158). Once the agarose polymerized, 200 µl of spinal cord medium were added to the drop and live imaging was started.

Live imaging recordings were carried out with an Olympus IX83 inverted microscope equipped with a spinning disk unit (CSU-X1 10’000 rpm, Yokogawa). Cultured spinal cords were kept at 37°C with 5% CO_2_ and 95% air in a PeCon cell vivo chamber. Temperature and CO_2_-levels were controlled by the cell vivo temperature controller and the CO_2_ controller units (PeCon). Spinal cords were incubated for at least 30 min before imaging was started. We acquired 18-35 planes (1.5 µm spacing) of 2×2 binned z-stack images every 15 min for 24 hours with a 20x air objective (UPLSAPO 20x/0.75, Olympus) and an Orca-Flash 4.0 camera (Hamamatsu) with the help of Olympus CellSens Dimension 2.2 software. Z-stacks and maximum projections of Z-stack movies were evaluated and processed using Fiji/ImageJ (Schindelin et al., 2012). Temporally color-coded projections were generated using Fiji/ImageJ. Kymograph analysis of axons crossing the floor plate or exiting it was performed as previously described (Medioni et al., 2015) using a region of interest (ROI) selection, the re-slice function and the z-projection of the re-sliced results in Fiji/ImageJ, which allowed following pixel movements within the horizontal axis. The ROI in the floor plate was selected as a 120×20 µm^2^ rectangle and the one in the post-crossing segment was a rectangle of 175×104 µm^2^. Note that the post-crossing segment ROI was rotated by 90° before running the kymograph analysis. The MtrackJ plugin (Meijering et al., 2012) was used to virtually trace single tdTomato-positive dI1 axons crossing the floor plate. Only axons that enter, cross and exit the floor plate during the 24-hour imaging period were traced and quantified. Overlays of traced axons with GFP and brightfield channels were used to assess the time the axons needed to cross the floor plate. Videos of axons with the ‘corkscrew’ phenotype were generated and assembled from z-stacks that were 2D deconvolved (nearest neighbor) using the Olympus CellSens Dimension 2.2 and Fiji/Image J software, respectively.

## Acknowledgements

We thank Dr. Beat Kunz and Tiziana Flego for excellent technical assistance, Alexandra Moniz and John Darby Cole for help with glycosylation dependence experiment, and members of the lab for discussions and critical reading of the manuscript. This project was supported by the Swiss National Science Foundation and the NCCR Brain Plasticity and Repair (Center of Transgenesis Expertise).

## Supplementary Figures

**Supplementary Figure 1.**
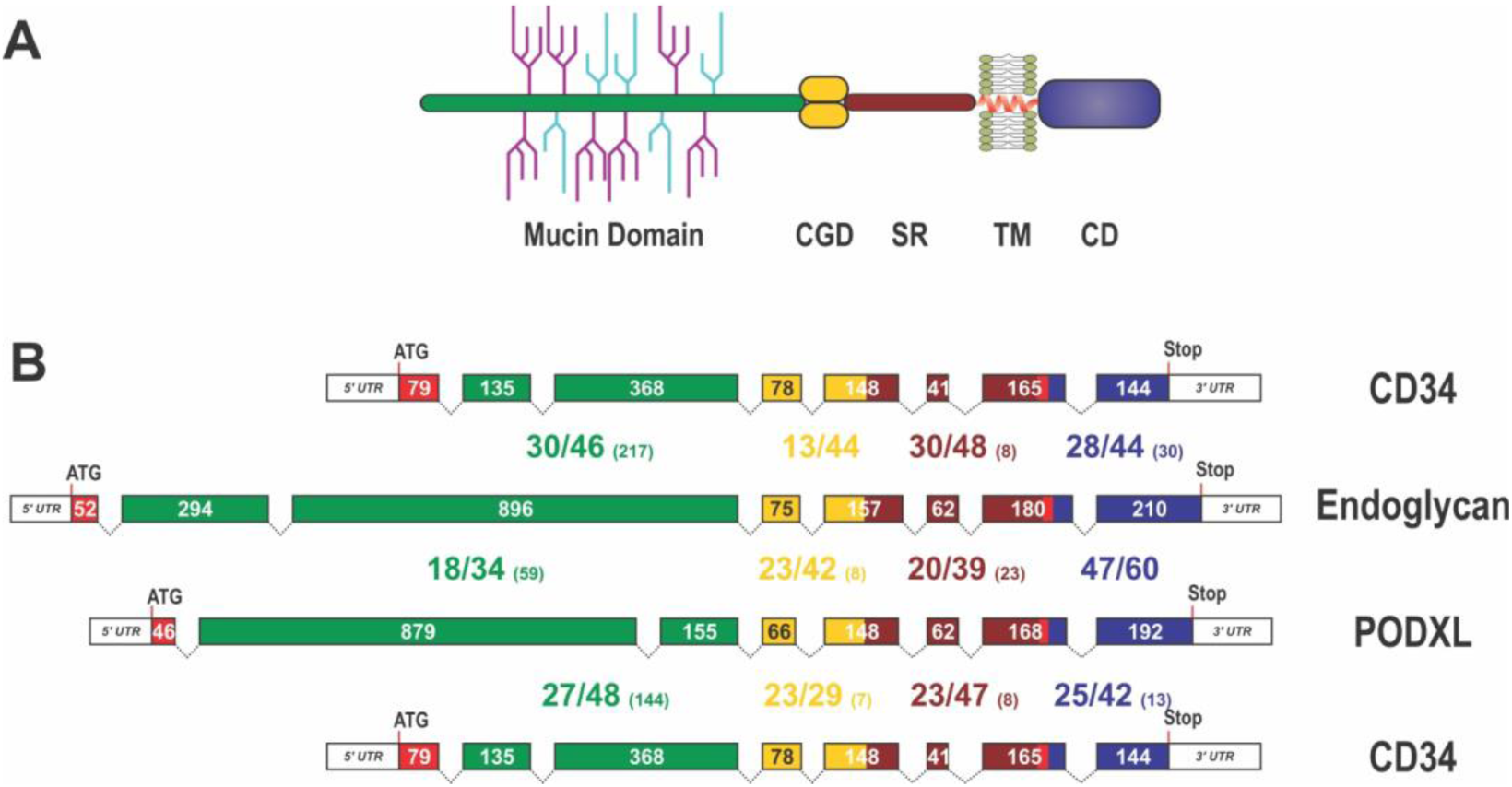
Domain organization, exon alignment and conservation of the CD34 family of sialomucins. (A) Schematic drawing of the domain organization of CD34 family members. They all contain an N-terminal, highly glycosylated, mucin domain (green), a cysteine-containing globular domain (CGD, yellow), a juxtamembrane stalk region (SR, brown), as well as a transmembrane alpha-helix (TM, red) and a cytoplasmic domain (CD, blue). O-linked glycosylation sites within the mucin domain are depicted in light blue, whereas further sialylated residues are symbolized in purple. N-linked glycosylation sites are not shown. Note, that the indicated glycosylation sites in this scheme are only symbolizing the extensive amount of glycosylation in sialomucins and are not representing the actual position of glycosylation. (B) Exon organization and domain conservation of sialomucins. Transcripts of CD34 family members are encoded by eight separate exons (colored boxes). While the length of exons coding for the cysteine containing globular domain (yellow), the juxtamembrane stalk region (brown), and the cytoplasmic domain (blue) are more or less conserved (exon sizes are given within the boxes), exons coding for the mucin domain (green) vary markedly in their length and organization. The translational start sites are highlighted by the ATG and the end of the coding sequences are indicated by the given Stop codon. Protein homology between the different chicken sialomucins is depicted by the large numbers between the exon pictograms. All domains were compared separately and the colors used indicate the corresponding domains. The first number indicates identical amino acids between the compared proteins, the second number represents conserved residues and the number in brackets designates the amount of gap positions within the alignment of the domains (eg. 28/44 (30)). The alignment was done using MUSCLE version 3.7. configured to the highest accuracy (Edgar, 2004). Single gap positions were scored with high penalties, whereas extensions of calculated gaps were less stringent. Using such parameters homologous regions of only distantly related sequences can be identified. Note, that within the mucin domain only some blocks, interspaced by sometimes large gap regions, are conserved between the different proteins.

**Supplementary Figure 2.**
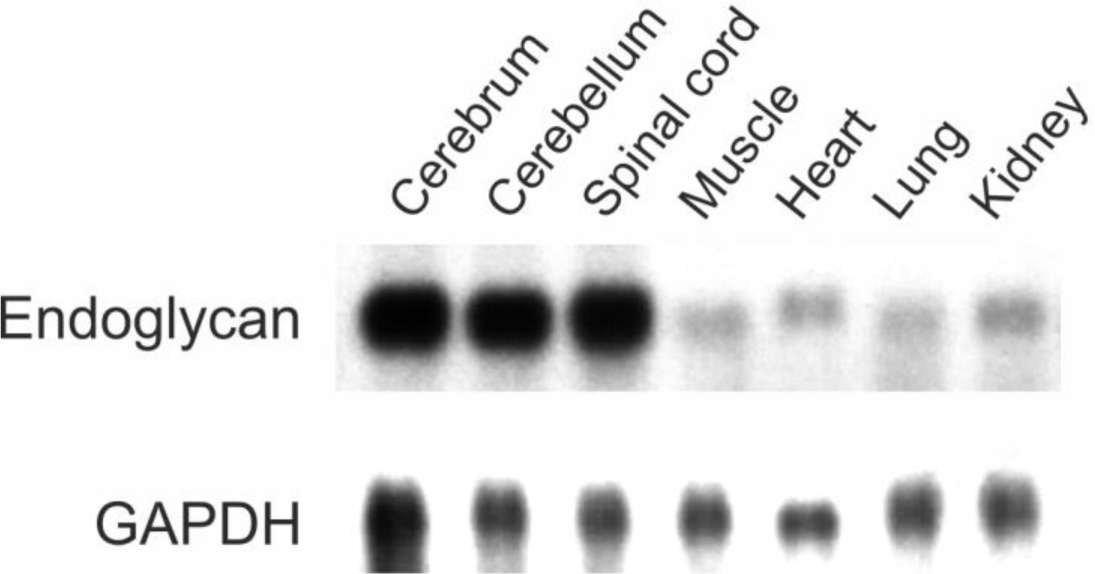
Endoglycan is mainly expressed in the developing nervous system. *Endoglycan* is expressed only at low levels in non-neuronal tissues. Northern blot analysis of tissues taken from HH38 chicken embryos revealed its high expression levels in the cerebrum (brain without cerebellum), the cerebellum, and the spinal cord. Only low levels were found in muscle, heart, lung, and kidney. GAPDH was used as a loading control.

**Supplementary Figure 3.**
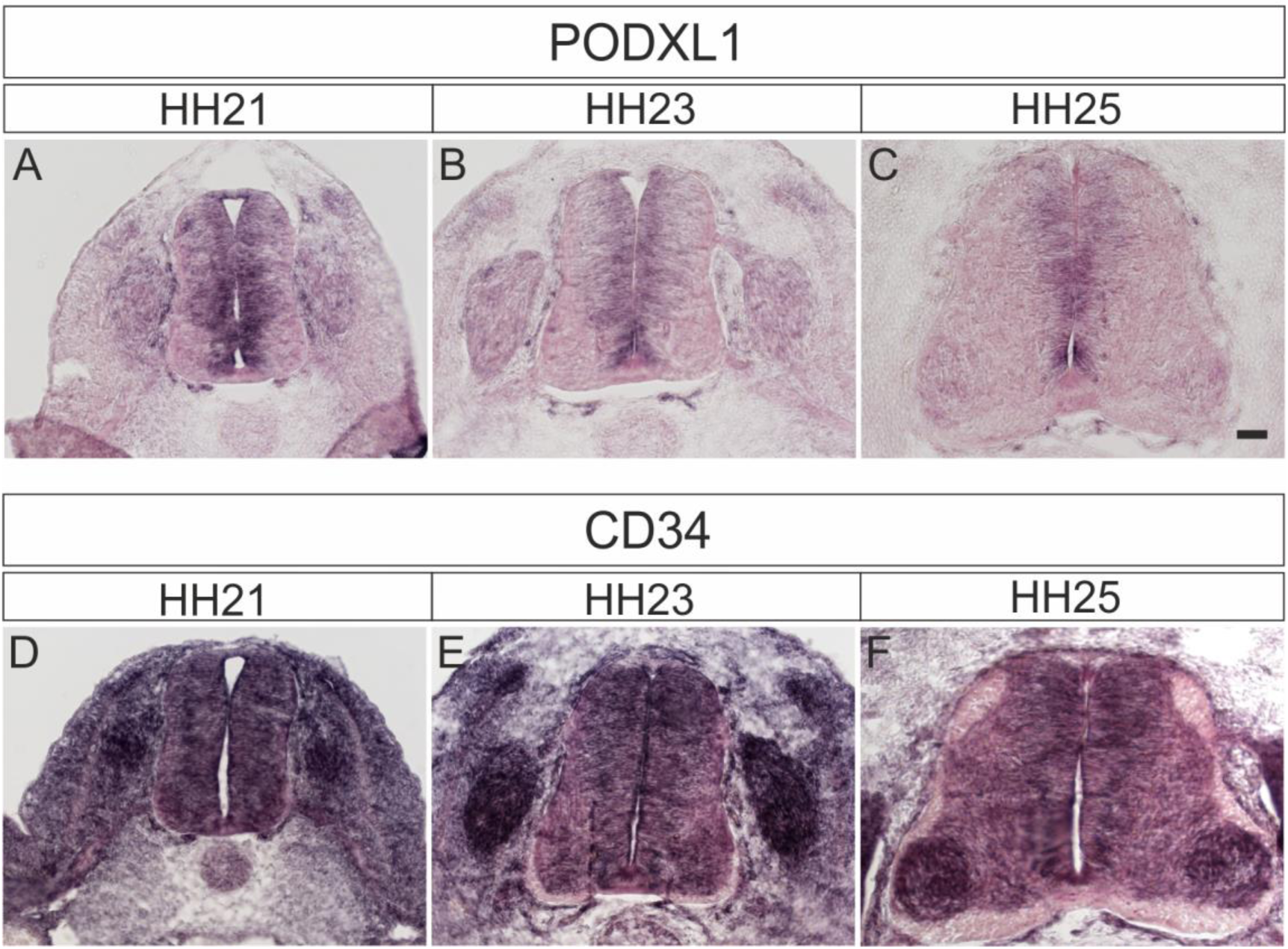
Podocalyxin and CD34 are expressed in the developing spinal cord during midline crossing of dI1 commissural axons. Podocalyxin1 expression was mainly found in the ventricular zone (A-C). At HH21, expression was found in precursors of dorsal interneurons (A). Expression was also seen in the dorsal root ganglia (DRG). Expression of CD34 (D-F) was more ubiquitous with highest levels in DRG and increasing levels in motoneurons at HH25 (F).

**Supplementary Figure 4.**
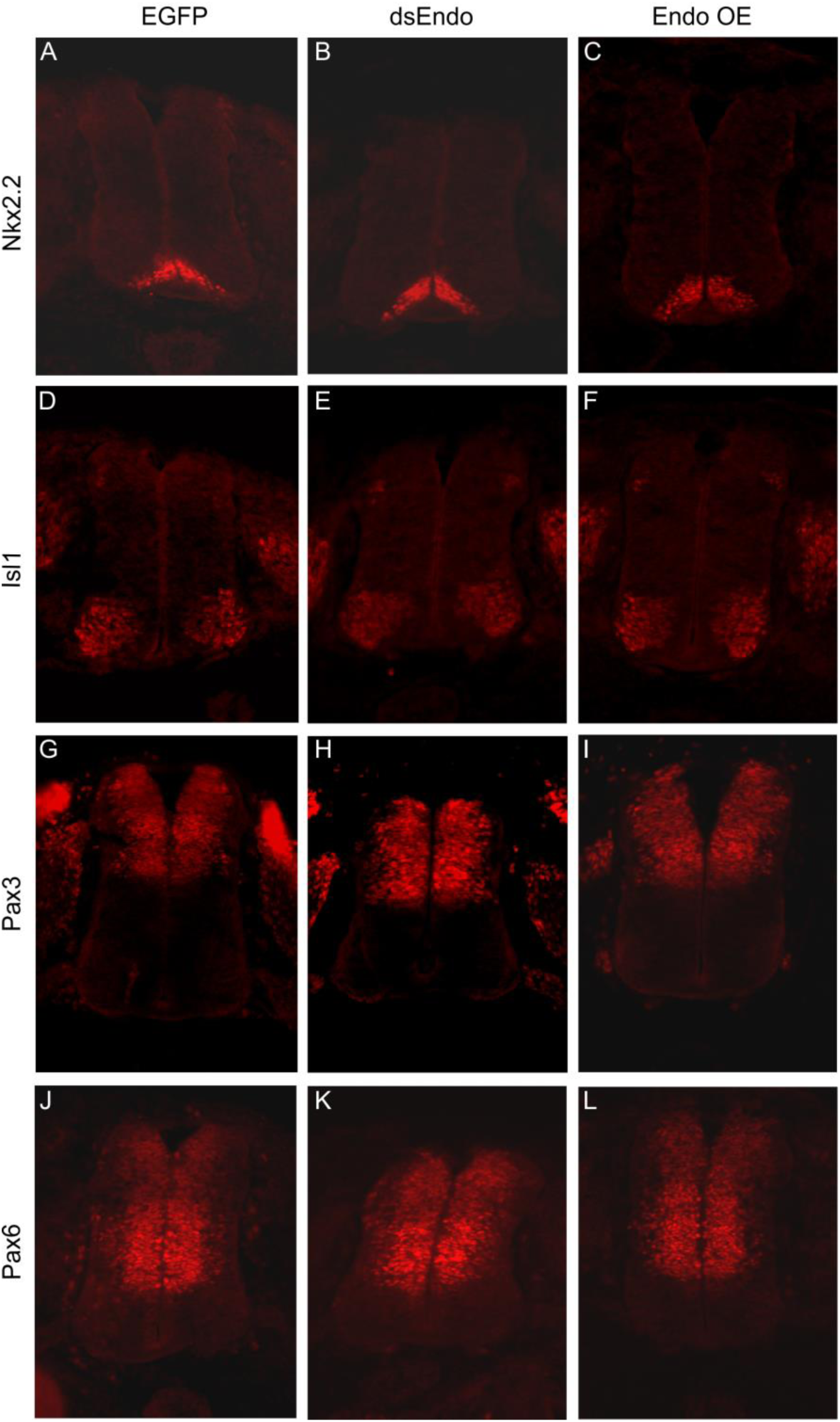
Downregulation or overexpression of Endoglycan does not affect spinal cord patterning. To check whether the effect of Endoglycan perturbation on axonal pathfinding was indirect due to changes in spinal cord patterning, we used a series of antibodies to stain sections taken from control embryos expressing GFP (A,D,G,J), embryos electroporated with dsEndo (B,E,H,K), or embryos overexpressing Endoglycan (C,F,I,L). We found no evidence for aberrant patterning, when we compared sections stained with Nkx2.2 (A-C), Islet1 (D-F), Pax3 (G-I), or Pax6 (J-L).

**Supplementary Figure 5.**
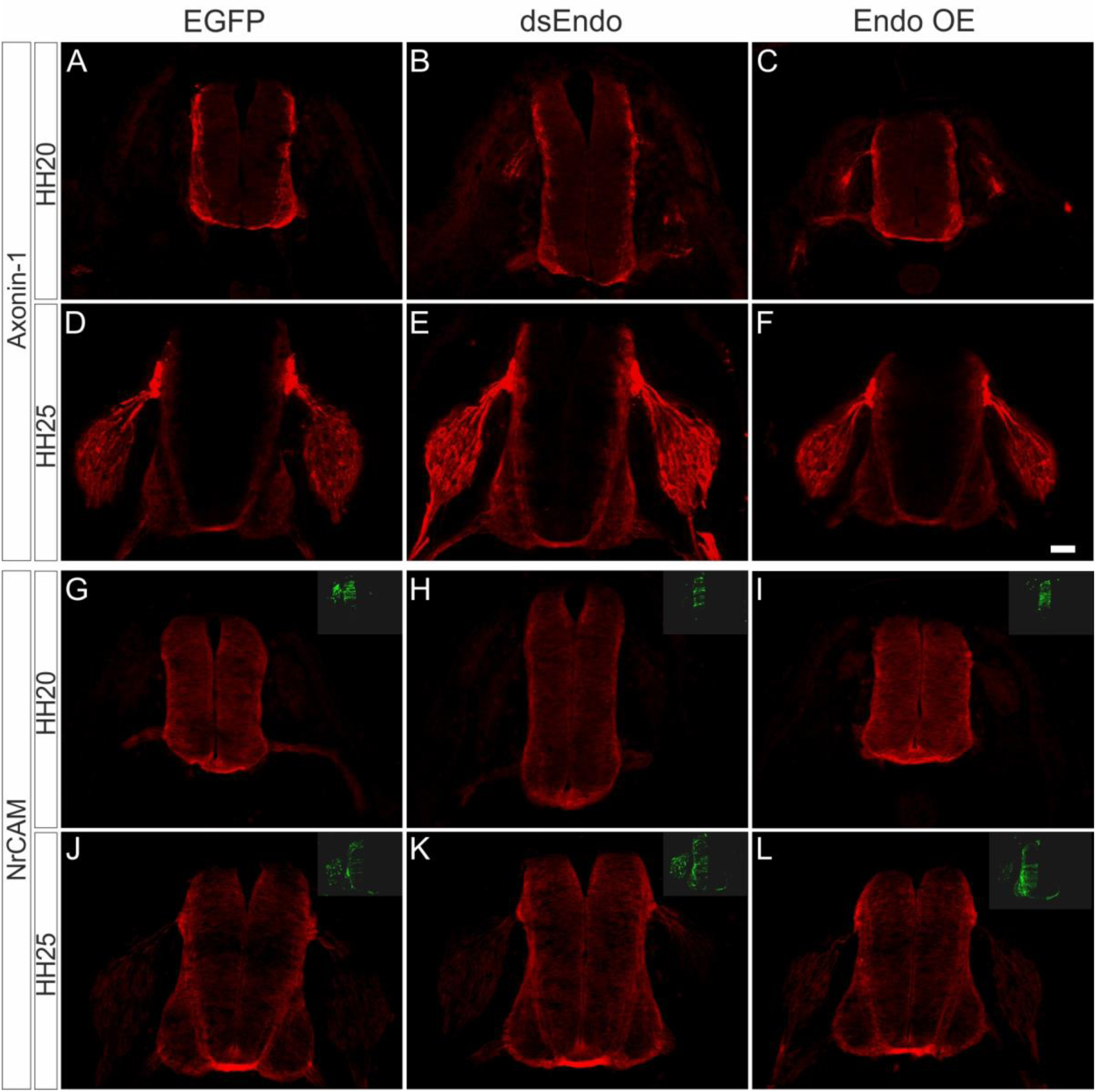
The errors in commissural axon pathfinding seen after perturbation of Endoglycan levels are not due to changes in the expression of known guidance cues for dI1 axons. To exclude that the changes in axonal pathfinding seen after silencing or overexpression of Endoglycan were explained by an effect on the expression of known axon guidance cues for dI1 axons, Axonin-1/Contactin-2 (A-F) or NrCAM (G-L), we compared sections taken from control embryos electroporated with a plasmid encoding GFP (A,D,G,J), embryos electroporated with dsEndo (B,E,H,K), or embryos overexpressing Endoglycan (C,F,I,L). We found no differences in expression of Axonin-1 and NrCAM. We compared sections taken from embryos sacrificed at HH20 (A-C, G-I) and HH25 (D-F, J-L). Bar: 50 µm.

**Supplementary Figure 6.**
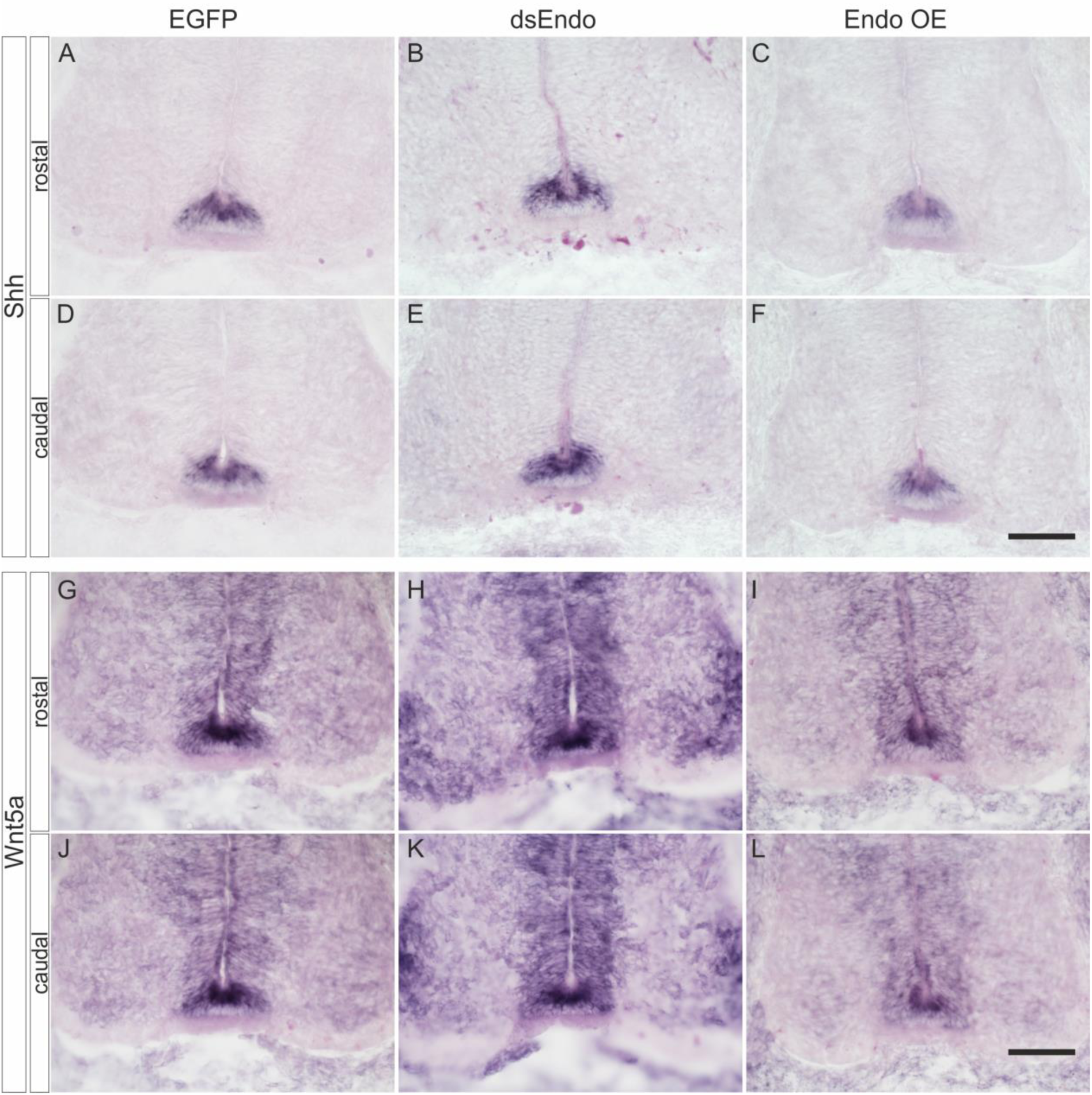
Perturbation of Endoglycan expression does not affect guidance of post-crossing commissural axons indirectly by changing Shh or Wnt5a expression. We did not find any changes in the expression of *Shh* (A-F) or *Wnt5a* (G-L) compared to control embryos expressing EGFP (A,D,G,J) after silencing *Endoglycan* (B,E,H,K) or after overexpression of *Endoglycan* (C,F,I,L). *Shh* was found at higher levels in the caudal compared to the rostral floor plate (A-F), as reported previously (Bourikas et al., 2005). *Wnt5a* levels did not differ between rostral and caudal sections taken from the lumbar part of the spinal cord, as reported earlier (Domanitskaya et al., 2010).

**Supplementary Figure 7.**
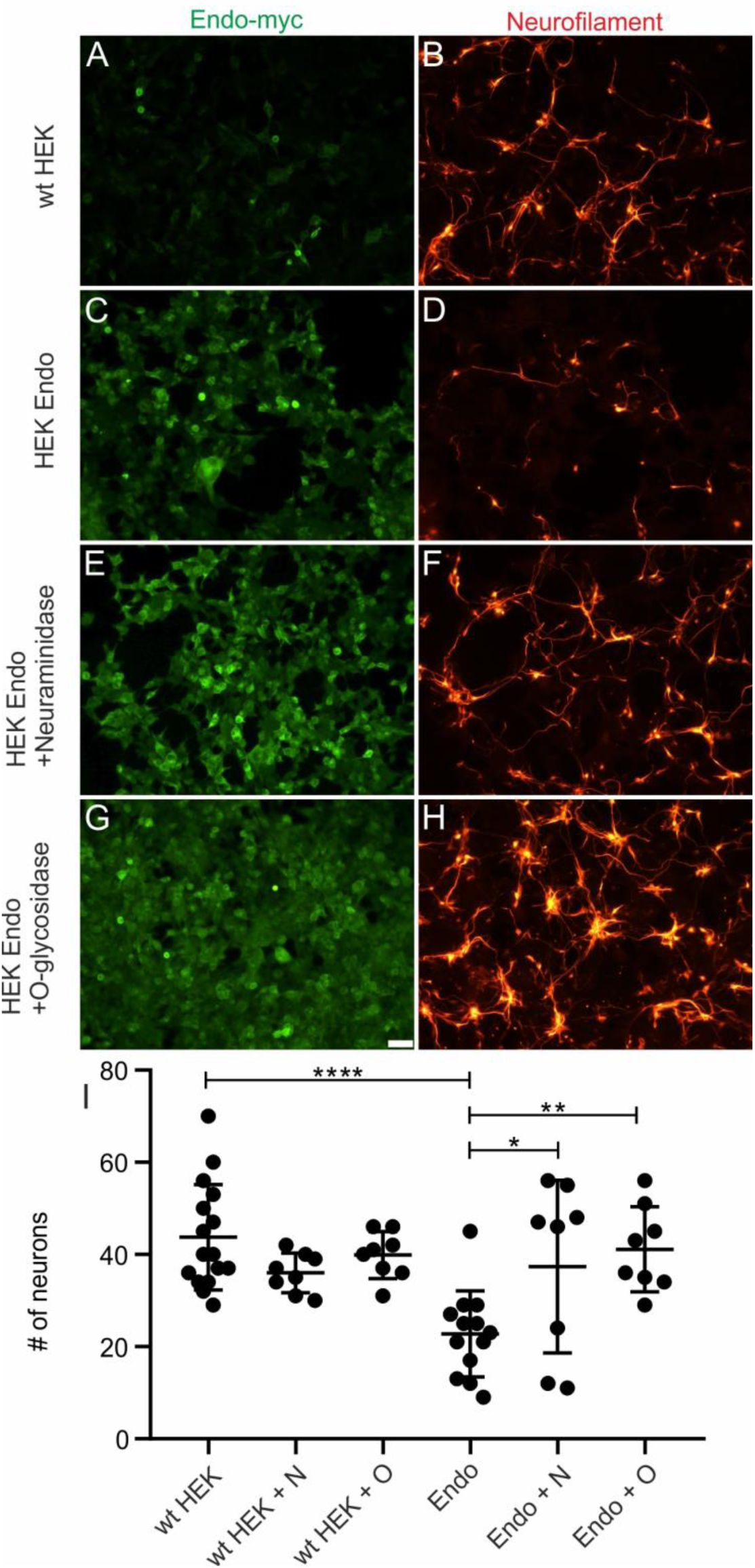
Post-translational modification of Endoglycan is required for its anti-adhesive effects. Control HEK cells (A-B) or Endoglycan-expressing HEK cells (C-H) were plated as carpet for motoneurons (stained with anti-neurofilament antibodies in B,D,F,H). Before adding the motoneurons, HEK cells were either left untreated (A-D), or treated with neuraminidase (E,F) or O-glycosidase (G,H) for 2 hours. Attached motoneurons were counted after 40 hours. Removal of sialic acid by neuraminidase (E,F) or O-glycosidase (G,H) abolished the anti-adhesive effect of Endoglycan (C,D). Quantification of the number of attached motoneurons under the different conditions is shown in (I).

**Supplementary Figure 8.**
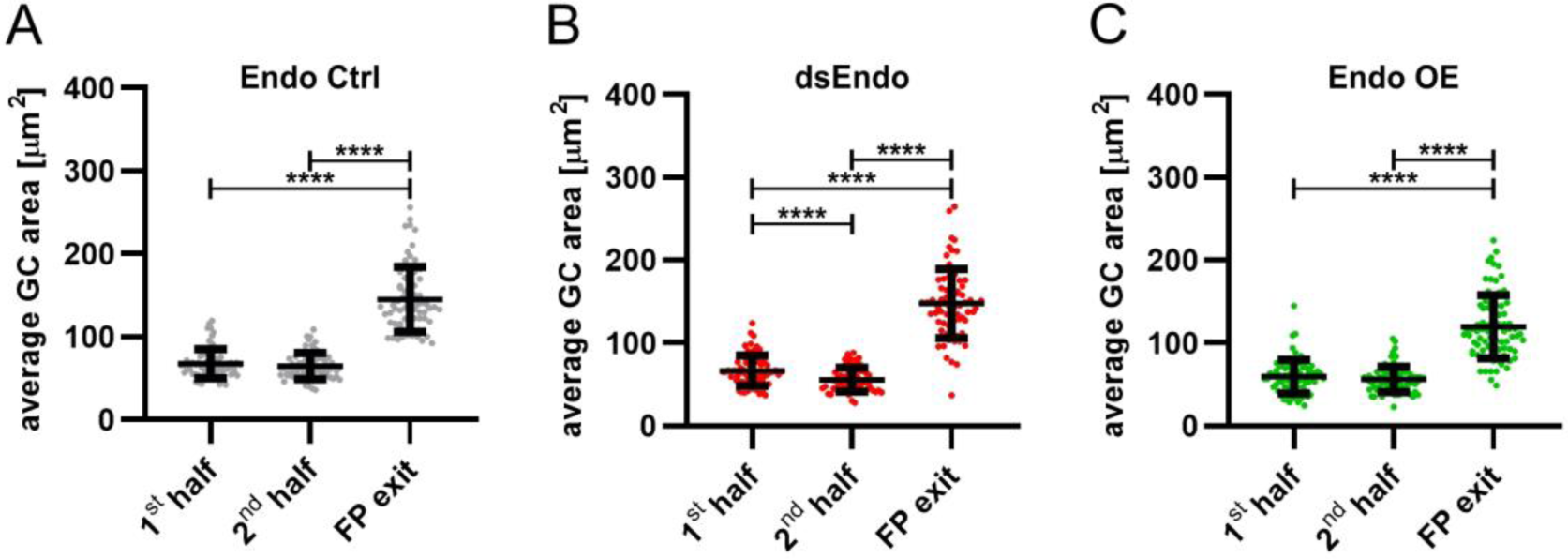
Growth cone size is enlarged at the floor-plate exit site. Data at the single axon level extracted from 24h time-lapse recordings of dI1 axons crossing the floor plate. The average growth cone area was measured in the first half, the second half and at the exit site of the floor plate for each condition. (A) No difference in the area was detected between growth cones in the first half and the second half of the floor plate in controls. However, growth cones were found to be much enlarged at the exit site compared to when they were in the floor plate (Friedman test with Dunn’s multiple-comparisons test). (B) Unilateral down regulation of Endoglycan induced a significant decrease in the average growth cone area in the second part of the floor plate compared to the first part. However, dI1 growth cones still got much larger when exiting the floor plate compared to when they were located in the floor plate (one-way RM ANOVA with Sidak’s multiple-comparisons test). (C) After unilateral overexpression of Endoglycan, no difference in growth cone area was detected between the first and the second half of the floor plate. Like in all other conditions, they were found to be much enlarged at the exit site (Friedman test with Dunn’s multiple-comparisons test). Error bars represent standard deviation. p<0.0001 (****), p<0.001 (***), p<0.01 (**), p<0.05 (*) and p≥0.05 (ns) for all tests. See Table 2 for detailed results.

**Supplementary Figure 9.**
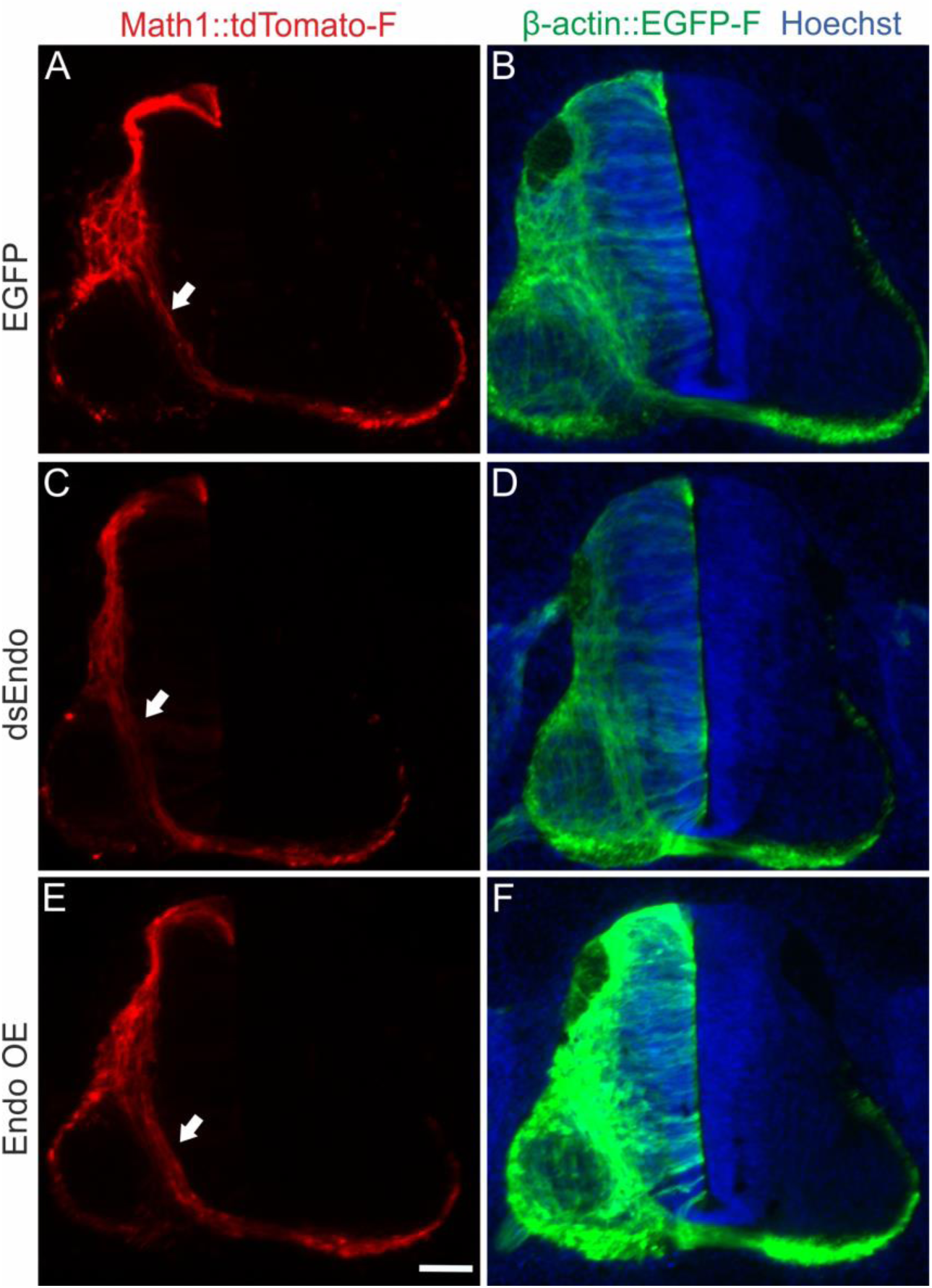
Endoglycan does not affect pre-crossing commissural axons. We found no differences in timing or trajectories of pre-crossing commissural axons labeled by the co-electroporation of Math1::tdTomato-F when we compared control embryos electroporated with an EGFP plasmid (A,B) with embryos electroporated with dsEndo (C,D) or embryos overexpressing Endoglycan (E,F).

**Supplementary Figure 10.**
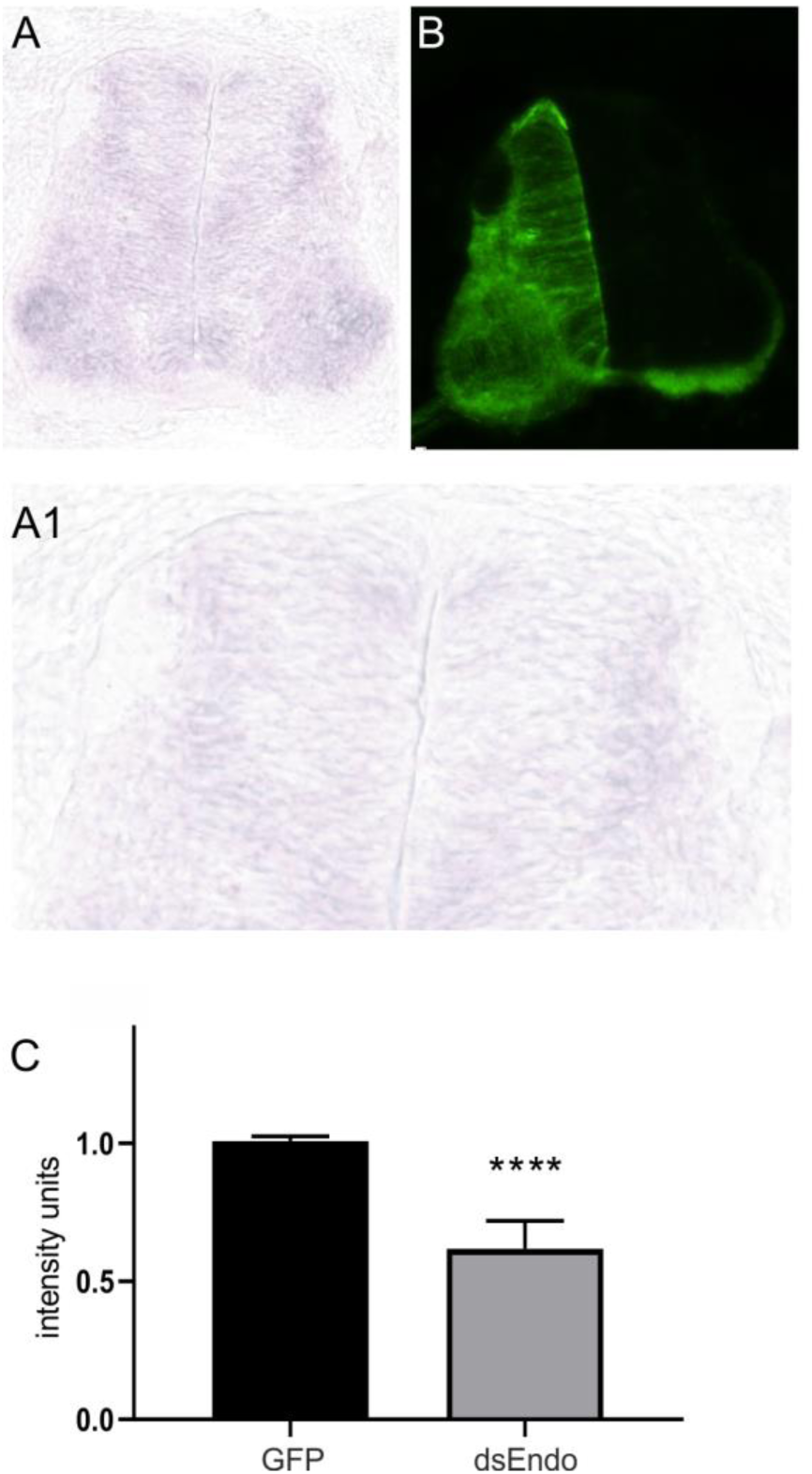
Electroporation of dsRNA derived from Endoglycan effectively downregulates Endoglycan mRNA. In the absence of antibodies detecting Endoglycan, we had to use in situ hybridization to quantify the efficiency of Endoglycan downregulation (A). When we compared expression levels of Endoglycan between the electroporated and the non-electroporated side of a HH25 spinal cord, we found reduced signal intensities on the electroporated side. (B) Co-electroporation of a plasmid encoding GFP together with dsEndo allows for the identification of the electroporated side. On average, electroporation of dsEndo reduced mRNA levels by 39%. Keep in mind that only about 50% of the cells in the electroporated area are taking up the dsRNA. Therefore, the reduction in Endoglycan is very efficient in the transfected cells. Three sections per embryo and three embryos per group were included in the analysis.

## Legends to supplementary movies

Supplementary Movie 1

24h time-lapse recordings of tdTomato-positive dI1 axons (shown in black) in control (Endo control), Endoglycan knockdown (dsEndo) and Endoglycan overexpression (Endo OE) conditions. Dashed lines represent floor plate boundaries. R, rostral; C, caudal.

Supplementary Movie 2

Representative examples of the floor plate crossing of tdTomato-positive dI1 axons (shown in black) taken from 24h time-lapse recordings from control (Endo control), Endoglycan knockdown (dsEndo) and Endoglycan overexpression (Endo OE) conditions. An arrowhead in each condition points at the migrating growth cone. Dashed lines represent floor-plate boundaries.

Supplementary Movie 3

Example of a 24h time-lapse recordings of tdTomato-positive dI1 axons (shown in black) showing guidance defects at the exit site of the floor plate in Endoglycan overexpression condition. Stalling growth cones are shown by arrowheads and caudally turning growth cones by arrows. Rostral is up. Dashed line represents the floor-plate exit site.

Supplementary Movie 4

The trajectory of single dI1 axons in the first half of the floor plate (electroporated half) in the absence of Endoglycan was aberrant and showed a ‘corkscrew’-like phenotype. Example of a tdTomato-positive dI1 axon taken from a 24h time-lapse recording of an ‘Endoglycan-knockdown’ spinal cord. Arrowheads show how the growth cone is migrating within the first half (electroporated half) of the floor plate and enter in contact with mislocated EGFP-positive cells (arrows) that induced a ‘corkscrew’ like morphology of the shaft. The 3D coronal rotation clearly shows that the axon and cells are located within the commissure (arrowhead).

Supplementary Movie 5

The trajectory of single dI1 axons in the first half of the floor plate (electroporated half) in the absence of Endoglycan was aberrant and showed a ‘corkscrew’-like phenotype. Example of tdTomato-farnesylated-positive dI1 axons taken from 24h time-lapse recordings of an ‘Endoglycan-knockdown’ spinal cord. Arrowheads show how two growth cones are migrating within the first half (electroporated half) of the floor plate. The first one made a loop within the floor-plate cell layer (arrowheads), as shown by the 3D coronal rotation. The second one was attracted towards mislocalized EGFP-positive cells (arrow), made contact with them and then carried out a U-turn toward them before continuing its migration in the direction of the midline. The 3D coronal view confirmed that the second growth cone during its U-turn and the mislocalized cells were located within the commissure (arrowhead).

